# Epigenetic therapy targets the 3D epigenome in endocrine-resistant breast cancer

**DOI:** 10.1101/2021.06.21.449340

**Authors:** Joanna Achinger-Kawecka, Clare Stirzaker, Neil Portman, Elyssa Campbell, Kee-Ming Chia, Qian Du, Geraldine Laven-Law, Shalima S. Nair, Aliza Yong, Ashleigh Wilkinson, Samuel Clifton, Heloise H. Milioli, Sarah Alexandrou, C. Elizabeth Caldon, Jenny Song, Amanda Khoury, Braydon Meyer, Julia M.W. Gee, Anthony Schmitt, Emily S. Wong, Theresa E. Hickey, Elgene Lim, Susan J. Clark

**Affiliations:** Epigenetics Research Laboratory, Cellular Science Pillar, The Garvan Institute of Medical Research, Sydney, New South Wales, Australia; St Vincent’s Clinical School, UNSW Sydney, New South Wales, Australia; Connie Johnson Breast Cancer Research Laboratory, The Garvan Institute of Medical Research, Sydney, New South Wales, Australia; Dame Roma Mitchell Cancer Research Laboratories, Adelaide Medical School, University of Adelaide, Adelaide, South Australia, Australia; Breast Cancer Molecular Pharmacology Group, School of Pharmacy and Pharmaceutical Sciences, Cardiff University, Wales, CF10 3NB, UK; Invasion and Metastasis Laboratory, The Garvan Institute of Medical Research, Sydney, New South Wales, Australia; Arima Genomics, Inc., San Diego, California, USA; Victor Chang Cardiac Institute, Sydney, New South Wales, Australia; School of Biotechnology and Biomolecular Sciences, UNSW Sydney, New South Wales, Australia

**Keywords:** 3D genome architecture, DNA methylation, Decitabine, Hi-C, epigenetic therapy, ER+ breast cancer, endocrine-resistance, transcription factors, enhancers

## Abstract

Three-dimensional (3D) epigenome remodelling is an important mechanism of gene deregulation in cancer. However, its potential as a target to overcome therapy resistance remains largely unaddressed.

Here we show that FDA-approved epigenetic therapy Decitabine (5-Aza-mC) suppresses tumour growth in preclinical metastatic ER+ breast tumour xenograft models. Decitabine-induced genome-wide DNA hypomethylation results in large-scale 3D epigenome deregulation, including de-compaction of higher order chromatin structure and loss of topologically associated domain boundary insulation. Significant DNA hypomethylation at ER-enhancer elements was associated with gain in ER binding, creation of ectopic 3D enhancer-promoter interactions and concordant activation of ER-mediated transcription pathways. Importantly long-term withdrawal of epigenetic therapy partially restores methylation at ER-enhancer elements, resulting in loss of ectopic 3D enhancer-promoter interactions and associated gene repression.

Our study illustrates how epigenetic therapy has potential to target ER+ endocrine-resistant breast cancer by DNA methylation-dependent rewiring of 3D chromatin interactions associated with suppression of tumour growth.

## Main

Around 70% of breast cancers are driven by the Estrogen Receptor-alpha (ER). ER is a critical ligand-activated transcription factor that controls breast cancer cell proliferation and tumour growth upon exposure to estrogenic hormones^1^. Drugs that target ER pathways are highly effective in the treatment of ER+ breast cancer^2,3^, however *de novo* or acquired resistance to these agents (endocrine resistance) affects a large proportion (>30%) of patients and is the major cause of breast cancer mortality. Endocrine resistance has previously been shown to be associated with epigenetic alterations, including DNA methylation, chromatin accessibility, histone modifications and binding of different transcription factors^4,5^. In particular, differential ER transcription factor binding leads to altered expression of estrogen-responsive genes in endocrine-resistant breast cancer^1^ and is associated with clinical response to endocrine therapies^6,7^.

Epigenetic alterations also influence the three-dimensional (3D) genome architecture, from the local level of chromatin interactions to the higher level organisation of topological associated domains (TADs) and chromosome compartments^8^. Alterations to the 3D genome architecture have been described in a number of different cancers, including prostate cancer^9,10^, breast cancer^11–15^, gliomas^16^, and several hematologic cancers^17,18^. Although cancer cells maintain the general pattern of 3D genome folding, distinctive structural changes have been described in cancer genomes at all of levels of 3D organisation^19^. We and others recently reported that 3D genome structure is also disrupted in endocrine-resistant ER+ breast cancer cells^14,15,20^, notably through long-range chromatin changes at ER-enhancer binding sites that are DNA hypermethylated in resistant cells^14^.

DNA demethylating agents such as Decitabine (5-aza-2’-deoxycytidine, DAC) have emerged as a promising therapeutic strategy for treating various cancers^21^. Decitabine is approved by many international regulatory agencies, including the US FDA and the European Commission (EC), for treating haematological cancers^21^. In solid cancers (including colorectal and ovarian cancer), Decitabine has been shown to demethylate regulatory regions that result in re-activation of tumour suppressor genes^22,23^. Additionally, treatment with DNA de-methylating agents has been shown to stimulate immune response pathways in cancer cells through increased transcription of DNA repeat elements, which induces a viral mimicry response^24–26^. However, the direct effect of these epigenetic drugs on the tumour cells, including epigenome and 3D genome structure, remains largely unexplored, especially in clinically relevant patient-derived model systems or clinical samples.

To elucidate the mechanism of epigenetic therapy with Decitabine, we assessed the molecular consequences of treatment on DNA methylation, 3D genome architecture and transcriptional programs in endocrine-resistant ER+ patient-derived xenograft (PDX) models. Our data revealed that Decitabine treatment inhibited tumour growth and resulted in DNA hypomethylation that was associated with 3D epigenome remodelling, gain in ER binding and activation of ER-mediated transcription, highlighting the potential of epigenetic therapy in treatment of ER+ endocrine-resistant breast cancer.

## Results

### Low-dose Decitabine inhibits tumour growth and decreases cell proliferation

To study the efficacy of epigenetic therapy in the context of endocrine-resistant ER+ breast cancer and to establish its impact on the 3D genome and epigenome, we used two different patient-derived xenograft (PDX) models (Gar15-13, HCI-005) (Fig. 1a) (see Methods). Gar15-13 and HCI-005 PDXs were derived from the metastases of ER+ patients who had disease progression following one or more lines of endocrine therapy and have been used for several pre-clinical studies^27–31^.

**Fig. 1.**
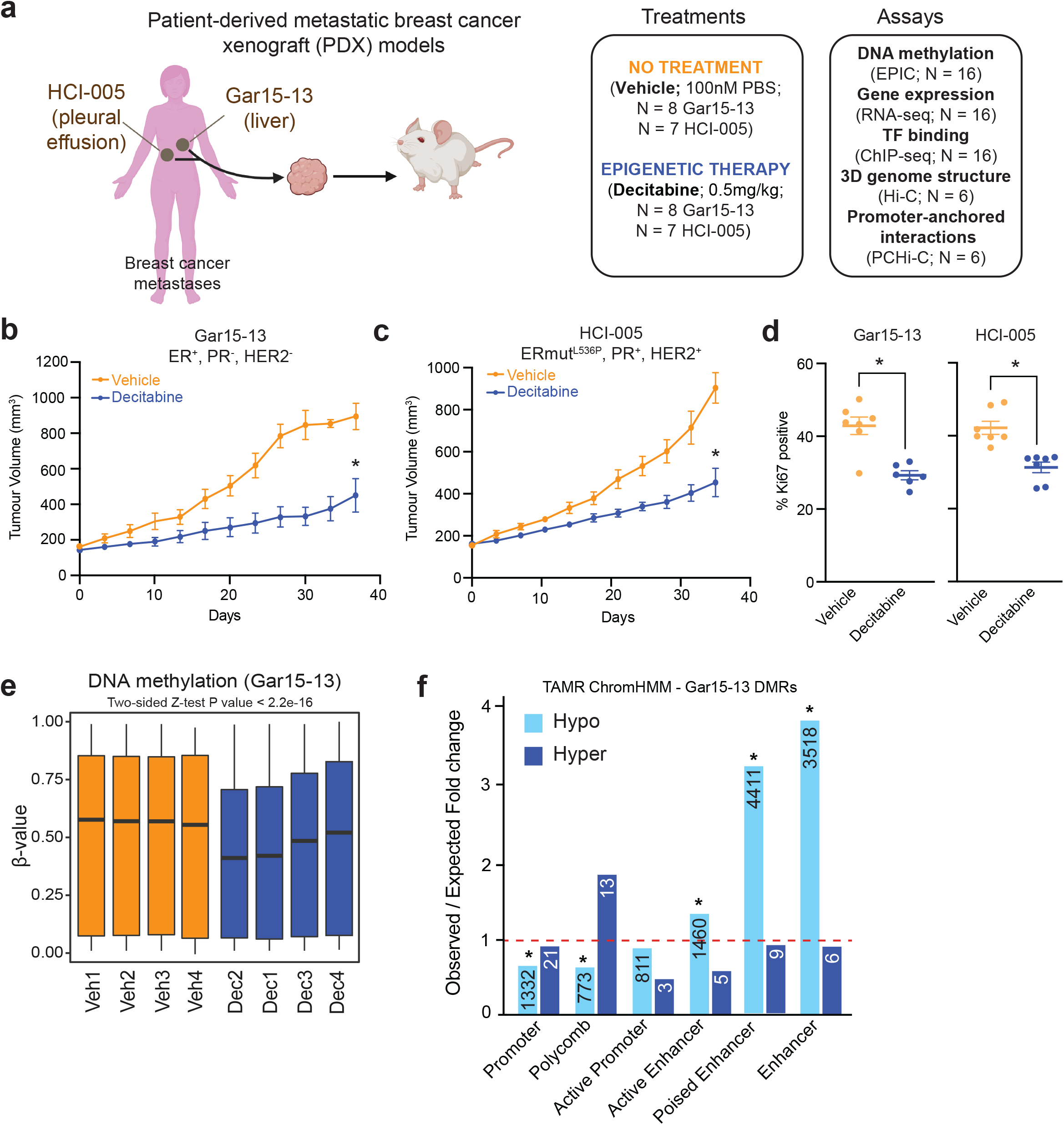
Decitabine inhibits tumour growth and induces widespread DNA hypomethylation. **(a)** Schematic of models and treatments used in the study and summary of genome-wide assays applied to PDX tumours at end-point. **(b)** Growth of treatment naïve (Vehicle) (100nM PBS, n = 7) and Decitabine-treated (0.5mg/kg, n = 7) Gar15-13 PDX tumours for 35 days or until ethical endpoint. All data are mean ± SEM. *** P value < 0.001. **(c)** Growth of treatment naïve (Vehicle) (100nM PBS, n = 8), and Decitabine-treated (0.5mg/kg, n = 7) HCI-005 PDX tumours for 35 days or until ethical endpoint. All data are mean ± SEM. * *P* value < 0.001. **(d)** Quantification of the proliferation marker Ki-67 at endpoint in Gar15-13 and HCI-005 tumours. Ki-67 proportions from all replicates per treatment were compared by two-tailed T test. * *P* value < 0.001. **(e)** Boxplots showing the distribution of DNA methylation profiles for Gar15-13 Vehicle and Decitabine-treated tumours for all EPIC probes across the genome. Black line indicates median ± SD. **(f)** Bar plot showing the association of differentially methylated probes in Gar15-13 Decitabine treatment compared to Vehicle across different regulatory regions of the genome as determined by TAMR ChromHMM annotation. Observed over expected fold change enrichment shown. * *P* value < 0.001. The numbers of hypo/hypermethylated probes located within each specific region are presented in the respective column.

Using a low, well-tolerated and non-cytotoxic dose (0.5mg/kg; Extended Data Fig. 1a) of Decitabine, we first interrogated the anti-cancer effect of epigenetic therapy on tumour growth. Following tumour implantation and initial period of growth (to a volume of 150-200 mm^3^), mice were randomized to twice-weekly injections of PBS (Vehicle) or 0.5mg/kg Decitabine. Treatment continued with twice-weekly measurement of tumour volume for 35 days or until tumour volume exceeded 1000 mm^3^. At endpoint mice were sacrificed and tumour material collected for analysis. In both Gar15-13 and HCI-005 PDX models, Decitabine treatment elicited a strong growth inhibitory response (Fig. 1b, c) and a significant reduction in the proliferative index as measured by Ki-67 positivity at endpoint (Fig. 1d).

### Decitabine induces widespread DNA hypomethylation

To directly identify the epigenetic reprogramming events induced by Decitabine treatment, we first analysed genome-wide DNA methylation data generated using Infinium EPIC Methylation arrays. EPIC arrays were performed on four biological replicates of Vehicle and Decitabine-treated PDX tumours at end-point. All Decitabine-treated tumours exhibited genome-wide DNA methylation loss (Fig. 1e and Extended Data Fig. 1b, c, *P* value < 0.0001, two-tailed Mann-Whitney test), with Gar15-13 showing more hypomethylation compared to HCI-005 (Fig. 1e and Extended Data Fig. 1d) (average methylation difference 14.55% and 8.74%, respectively). To quantify the extent of DNA methylation loss, we identified differentially methylated probes (DMPs) and differentially methylated regions (DMRs) between Vehicle and Decitabine-treated Gar15-13 and HCI-005 tumours (Supplementary Table 1). Hypomethylated DMPs in both PDX models were mainly located at non-coding genomic regions (introns and intergenic) (Extended Data Fig. 1e, f) and significantly enriched at putative enhancer regions (Fig. 1f and Extended Data Fig. 1g) (*P* < 0.001, permutation test). In agreement, we observed extensive DNA hypomethylation at enhancers in Gar15-13 tumours (approx. 18.38% change in median DNA methylation; Extended Data Fig. 1h, *P* value < 0.0001, two-tailed Mann-Whitney test) and HCI-005 tumours (approx. 9.24% change in median DNA methylation; Extended Data Fig. 1i, *P* value < 0.0001, two-tailed Mann-Whitney test), while promoters were less demethylated (approx. 10.12% in Gar15-13 and 2.16% in HCI-005; Extended Data Fig. 1j-k, *P* value < 0.0001, two-tailed Mann-Whitney test).

We next evaluated genome-wide DNA methylation levels at different classes of transposable elements (TEs) using the REMP package^32^, including LTRs, LINE1 elements and Alu elements. We observed genome-wide loss of DNA methylation at all repeat element sub-groups (Extended Data Fig. 1l and Extended Data Fig. 1m) with ~12% loss of DNA methylation in Decitabine treated tumours. Additionally, we observed TE expression alterations and activation of anti-viral signalling (Supplementary Note), as previously observed with Decitabine treatment in other cancers^24,25^. Notably, although all TE classes showed some degree of DNA methylation loss, the extent of DNA hypomethylation measured at repeat elements was less than those observed genome-wide and significantly less compared to DNA hypomethylation at enhancer regions (Extended Data Fig. 1h-i).

### Loss of DNA methylation results in de-compaction of 3D genome architecture and alters TAD boundary insulation

Distinctive structural changes have been described in cancer genomes at all levels of 3D organisation^19^. We were therefore motivated to study if Decitabine-induced genome-wide DNA hypomethylation also leads to global changes in 3D genome architecture. We analysed *in situ* Hi-C data (see Supplementary Note) corresponding to three biological replicates of Vehicle and Decitabine-treated tumours in Gar15-13 PDX. First, to detect open (active) and closed (silent) genomic compartments (A and B, respectively) we performed PCA analysis of the Hi-C data as described^33^. Comparison of eigenvalues between samples revealed differences between Vehicle and Decitabine tumours as shown by reduced correlation (Extended Data Fig. 2a). To further understand these differences, we directly compared the eigenvalues between Vehicle and Decitabine-treated tumours and observed that while most bins retained the same compartment status between the samples (either A to A or B to B), a large number of bins became more A-type in Decitabine as compared to Vehicle (B to A switch) (Fig. 2a). We next quantified this switching (see Methods) and identified 643 compartments (i.e. 2.28% of all compartments) that switched their assignment between A and B (Extended Data Fig. 2b). Around 64% of the changes induced by Decitabine involved compartment activation (B-type to A-type) (Fig. 2b). Notably, we observed significant DNA hypomethylation at B to A switches, while A to B switching regions maintained similar DNA methylation levels to Vehicle samples (Fig. 2c). In agreement with B-compartments becoming more de-methylated and active with Decitabine, using RNA-seq data (see Supplementary Table 4) we detected an overall increase in expression of genes located at regions that switched their assignment from B to A in Decitabine-treated tumours, while genes located at A to B switching compartments did not significantly change expression (Extended Data Fig. 2c). The newly activated compartments hosted 21 genes that decreased in expression and 87 genes that increased in expression with Decitabine (Extended Data Fig. 2d, e). Furthermore, we quantified A–A and B–B interaction frequencies in Decitabine and Vehicle-treated tumours and found decreased interaction strength between closed compartments (B-B interactions; *P* = 0.025, two-tailed Students t-test), increased contacts between active compartments (A-A interactions; *P* = 0.26, two-tailed Students t-test) and increased contacts between A-B compartments (*P* = 0.011, two-tailed Students t-test) (Fig. 2d, e and Extended Data Fig. 2f, g).

**Fig. 2.**
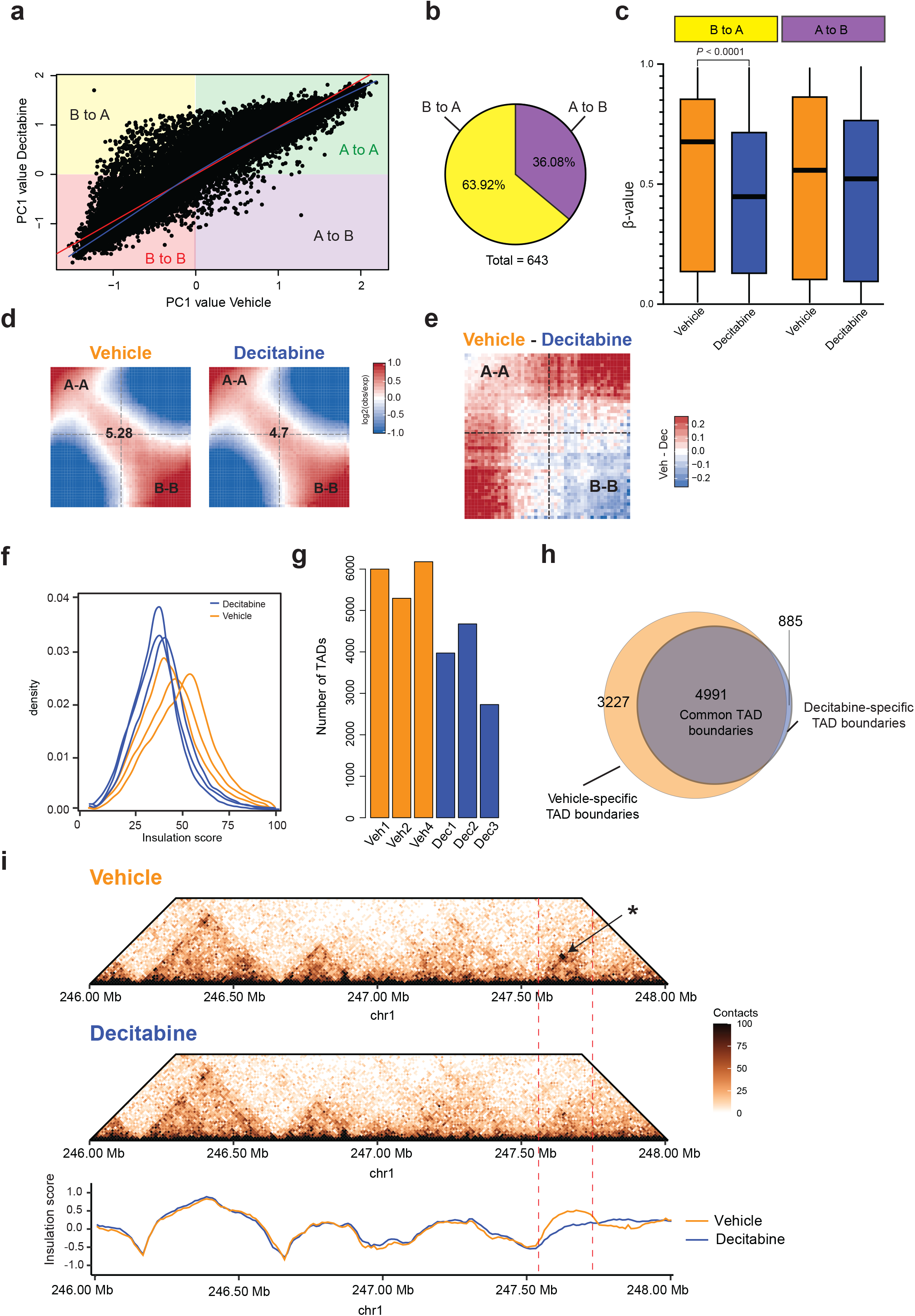
Loss of DNA methylation results in de-compaction of chromatin. **(a)** Scatterplot showing correlation between average eigenvalues per bin in Vehicle and Decitabine-treated Gar15-13 PDX tumours. **(b)** Pie chart showing distribution of different types of switching compartments (A to B; B to A) in Decitabine-treated tumours as compared to Vehicle. **(c)** Boxplot showing DNA methylation levels at compartment regions that switched their assignment from B to A and from A to B in Decitabine-treated (n = 4) and Vehicle (n = 4) PDX tumours. Black line indicates median ± SD. **(d)** Average contact enrichment (saddle plots) between pairs of 50Kb loci arranged by their PC1 eigenvector in Vehicle and Decitabine-treated tumours. Average data from n = 3 Hi-C replicates for Vehicle and Decitabine shown. The numbers at the centre of the heatmaps indicate compartment strength calculated as the log2 transformed ratio of (A–A + B–B)/ (A–B + B–A) using the mean values. **(e)** Difference in Hi-C data between Vehicle and Decitabine treatments. Saddle plots were calculated using the averaged PC1 obtained from Vehicle (n = 3) and Decitabine-treated (n = 3) tumours. **(f)** Density plot of insulation scores calculated in Vehicle and Decitabine-treated tumours. **(g)** Number of TADs identified in Vehicle (n = 3) and Decitabine-treated (n = 3) PDX tumours. **(h)** Venn diagram showing overlap between TAD boundaries identified in Vehicle and Decitabine-treated tumours. **(i)** Snapshot of region on chromosome 1, showing Vehicle (top panel) and Decitabine-treated tumour Hi-C matrixes (bottom panel), demonstrating loss of a TAD in Decitabine-treated samples (indicated with an arrow), concomitant with decreased insulation at that region. Merged Hi-C data from replicates (n = 3 each) shown at 10Kb resolution.

We next investigated the impact of Decitabine treatment on the organisation of topologically associated domains (TADs). We observed a significant decrease in the average insulation score in Decitabine as compared to Vehicle-treated tumours (~36.53 in Decitabine and ~46.74 in Vehicle) (Fig. 2f and Extended Data Fig. 2h, *P* value < 0.0001, two-tailed Mann-Whitney test), suggesting that loss of DNA methylation is associated with reduced segmentation into TADs. Consistent with loss of TAD boundaries and potential merging of TADs, the total number of TADs was decreased in Decitabine-treated samples (Fig. 2g) and their corresponding average domain size increased (Extended Data Fig. 2i; *P* = 0.0289, t-test). Analysis of differential TAD boundaries revealed a large percentage (43.2%) of Vehicle-specific boundaries, which were lost in Decitabine tumours (Fig. 2h) and characterised by decreased average insulation score in Decitabine tumours (Extended Data Fig. 2j). We exemplify one such region, where a TAD was lost in Decitabine-treated tumours, concomitant with loss of boundary insulation (Fig. 2i; further examples in Extended Data Fig. 2k-l).

Together, these results indicate that DNA hypomethylation induced by Decitabine treatment *in vivo* leads to significant de-compaction of 3D chromatin architecture, with reduced B-type compartments, increased interactions within A-type compartments and concomitant increase in regional gene expression. While most TADs maintain their structure after Decitabine treatment, their boundaries become less insulated suggesting increased intra-tumour heterogeneity in TAD structure and loss of some TAD boundaries at the regions of chromosomal compartment de-compaction.

### Loss of DNA methylation results in rewiring of 3D enhancer-promoter interactions

Since we observed altered higher-level 3D genome structure and significant hypomethylation at enhancer elements with Decitabine treatment (Extended Data Fig. 1h), we next asked if there were alterations in finer-scale 3D chromatin interactions. To gain insights into chromatin interactions at the level of individual promoters and enhancers, we investigated genome-wide promoter-anchored contacts in three Decitabine and three Vehicle-treated tumours using Promoter Capture Hi-C (PCHi-C) (Supplementary Note). Hi-C libraries were hybridised to custom-designed genomic restriction fragments covering 23,711 annotated gene promoters, achieving a clear increase in the coverage of promoter-anchored interactions as compared to Hi-C (Fig. 3a). To identify statistically significant interactions between promoters and other regulatory elements from the PCHi-C data, we used the CHiCAGO pipeline^34^. Promoter (bait) regions were significantly enriched for active promoters, poised promoters and active enhancers ChromHMM states in both Vehicle and Decitabine-treated tumours (Fig. 3b). Notably, putative enhancer other-end interacting regions (i.e. enhancer OEs; exemplified in Fig. 3a) showed significant differential enrichment, whereby active promoters were enriched in Vehicle tumours and enhancers were enriched in Decitabine-treated tumours, especially active enhancers (Fig. 3b).

**Fig. 3.**
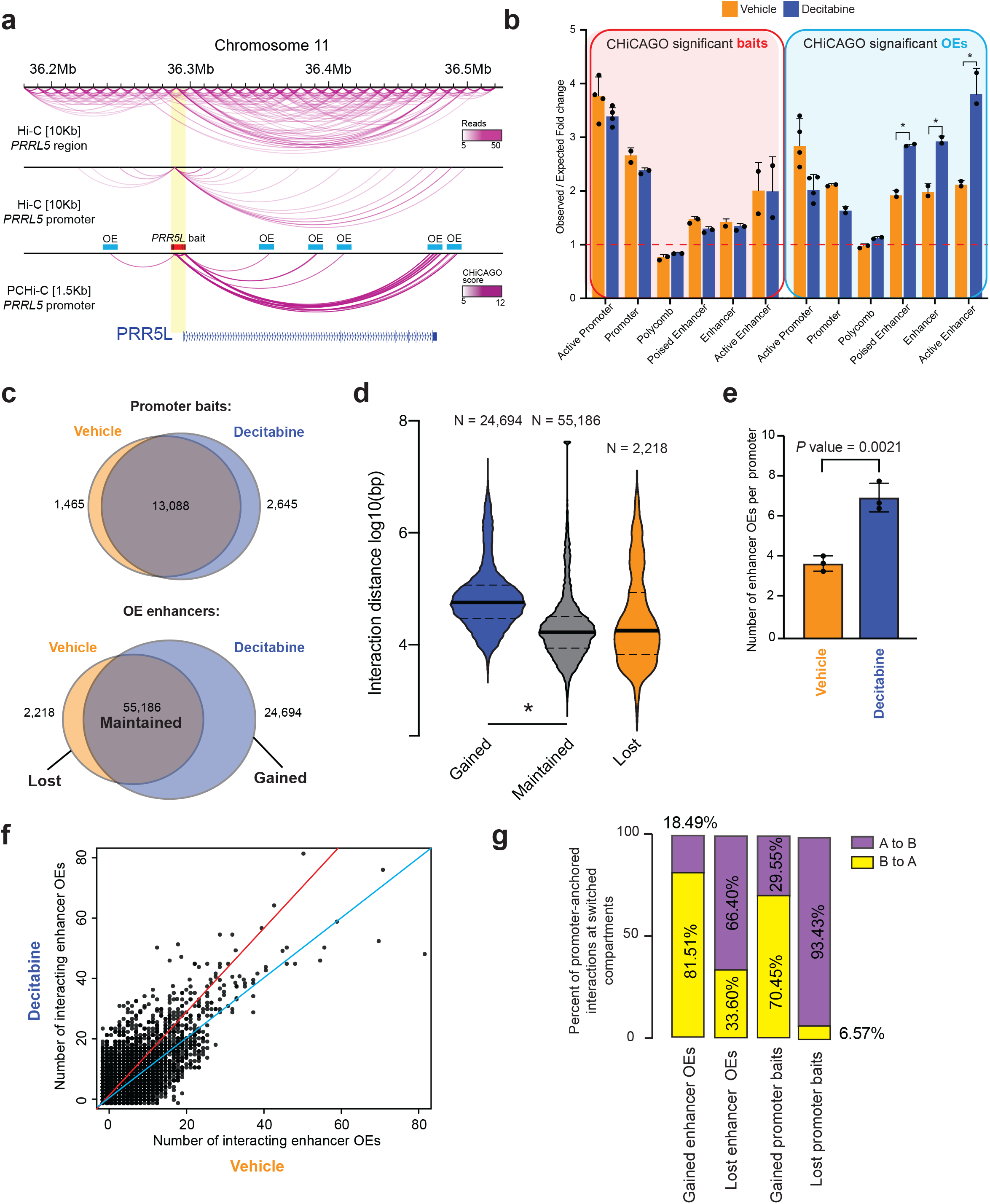
Loss of DNA methylation rewires 3D enhancer-promoter interactions. **(a)** Browser snapshot of interactions landscape at the *PRR5L* gene demonstrating increased coverage of promoter-anchored interactions in PCHi-C at 1.5Kb resolution compared to Hi-C at 10Kb resolution. Bait and other end (OE) regions marked for illustrative purposes. **(b)** ChromHMM (TAMR) annotation of CHiCAGO significant interaction bait (promoter) and other end (OEs) regions (putative enhancers) in Decitabine-treated and Vehicle samples (* *P* value < 0.001) **(c)** Overlap between promoter bait and other end (OE) enhancer regions for CHiCAGO significant interactions in Vehicle and Decitabine-treated tumours. Merged data across replicates shown. **(d)** Violin plots showing the log10 genomic distance of promoter interactions whose enhancer OEs are gained, maintained or lost following Decitabine treatment. *P* value < 0.0001, Wilcoxon test. Merged data across replicates shown. **(e)** Average number of enhancer OE interactions per promoter bait. Error bars indicate the interquartile range. *P* value Wilcoxon test. **(f)** Scatterplot showing number of enhancer OE interactions per promoter bait for each CHiCAGO significant promoter-anchored interaction in Vehicle and Decitabine-treated tumours. Merged data across replicates shown. **(g)** Percent of overlap of promoter baits and enhancer OEs that are either gained or lost in Decitabine with compartment that switch with Decitabine (A to B or B to A).

In order to directly identify differential promoter-anchored interactions, we integrated the results generated using Chicdiff^35^ pipeline with methods to intersect the promoter bait and enhancer OE regions for each interaction (see Methods). In total, we found 13,088 stable (no change) and 4,111 dynamic (gained or lost) contacts for promoters and 55,186 stable and 26,912 dynamic contacts for enhancer OEs (Fig. 3c). The majority of promoter regions were common between the Decitabine and Vehicle tumours. However, Decitabine treatment resulted in a large gain in the number of dynamic enhancer OEs, while only a small number of enhancer OEs were lost (24,694 gained and 2,218 lost in Decitabine) (Fig. 3c). Notably, interactions at gained enhancer OEs with Decitabine treatment were associated with longer interaction distances as compared to these that were at maintained or lost (Fig. 3d), consistent with an increased number of long-range interacting enhancers connecting to these promoters.

We next compared the total number of unique promoter and enhancer OE regions involved in interactions between Vehicle and Decitabine-treated tumours and found a significant increase in the number of enhancer OEs in Decitabine tumours while the number of interacting promoters stayed the same (Extended Data Fig. 3a, b). On average we detected 3.73 unique enhancer OEs per promoter in Vehicle samples and 7.06 unique enhancer OEs per promoter in Decitabine samples (Fig. 3e). To directly assess the gain in the number of interacting enhancers with Decitabine, we calculated the number of interacting enhancer OEs per each individual promoter and compared this number between Vehicle and Decitabine-treated tumours (Fig. 3f). We found that the majority of bait promoters involved in interactions in Vehicle tumours showed a large gain in the number of enhancer OEs they connect to in Decitabine-treated tumours, suggesting reprogramming of one-to-many enhancer-promoter interactions. Furthermore, we identified gained multi-way interactions that had, on average, significantly higher CHiCAGO scores in Decitabine-treated tumours compared to Vehicle (Wilcoxon *P* value < 0.0001) (Extended Data Fig. 3c), consistent with an overall increase in the total number of interactions with Decitabine treatment.

Given that the frequency of interactions is directly linked with higher-order compartment assignment, we next measured the overlap between dynamic promoter-anchored interactions and A/B compartment flipping. We found that the gain of interactions was associated with a shift from B compartment assignment toward compartment A in Decitabine-treated tumours (~76%) (Fig. 3g and Extended Data Fig. 3d), while a loss of interactions was associated with switch from A-type to B-type assignment (~80%). This was particularly pronounced at lost interactions involving promoter bait regions (>90% switched A to B) (Fig. 3g).

Together, these results support that 3D chromatin interactions are rewired following Decitabine-induced DNA methylation loss, leading to increased promoter-anchored interactions involving multiple enhancers connecting to gene promoters.

### Rewiring of 3D chromatin interactions aligns with altered transcription and gain in ER binding at ER-mediated enhancers

To examine the transcriptional consequences of Decitabine-induced rewiring of 3D chromatin interactions, we next analysed RNA-seq data corresponding to four replicates of Decitabine-treated and Vehicle Gar15-13 PDX tumours. Gene set enrichment analysis (GSEA)^36^ of all differentially expressed genes revealed that Decitabine treatment negatively correlated with gene signatures of cell proliferation and cell cycle (E2F targets, G2M checkpoint and Myc targets; for example *SMC3, TK1* and *KIF20A*) (Fig. 4a), consistent with supressed tumour growth and decreased cytological proliferation markers observed (Fig. 1d), as well as genes involved in viral mimicry response (see Supplementary Note), as described previously^25^. Surprisingly, Decitabine treatment also enriched for multiple hallmark genesets related to hormone signalling (estrogen response early and estrogen response late) (Fig. 4a) and upregulation of a significant proportion of genes belonging to the Hallmark Estrogen Response genesets (Extended Data Fig. 4a).

**Fig. 4.**
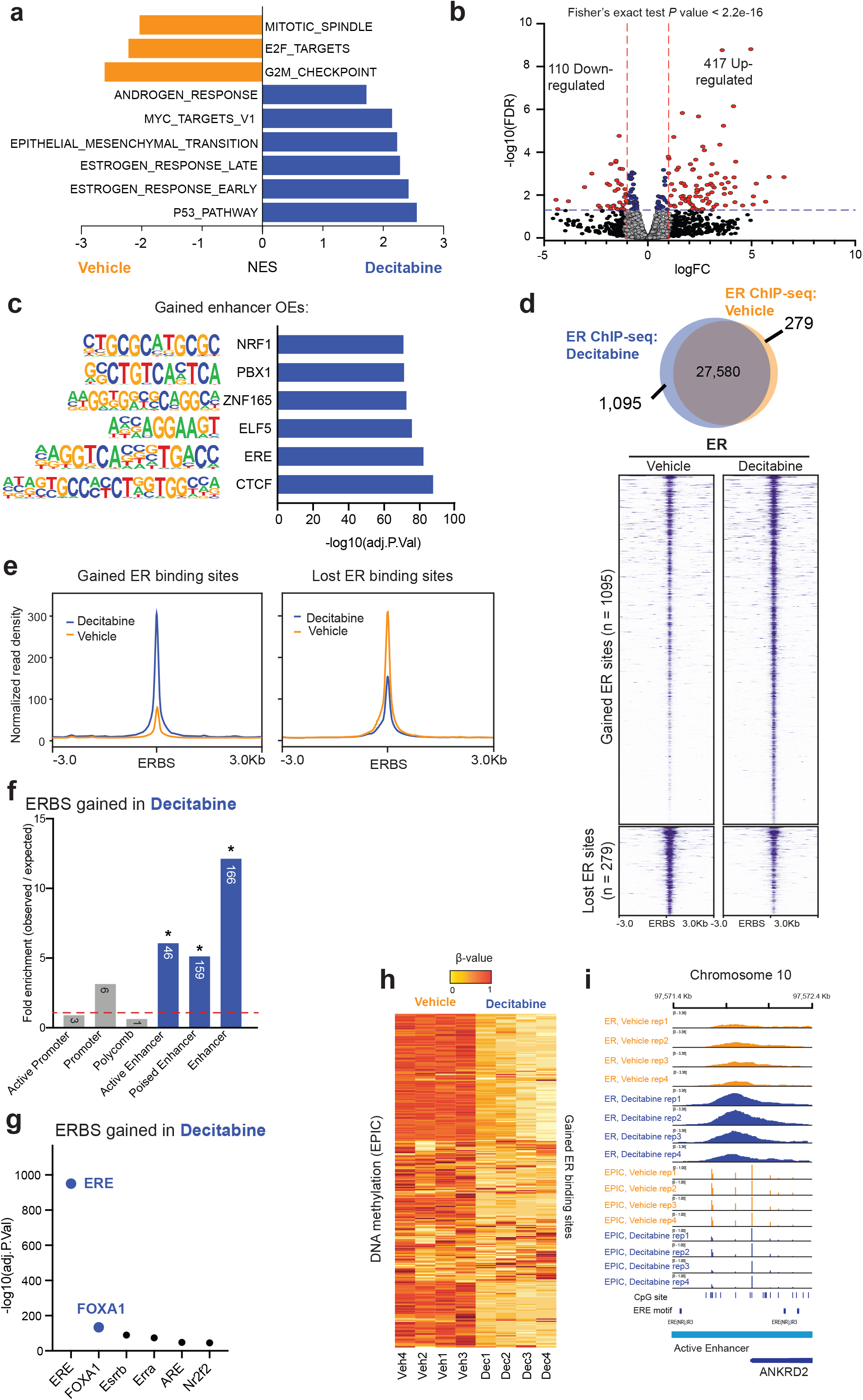
Rewired 3D chromatin interactions align with altered transcription. **(a)** Normalized enrichment scores (NES) for signature gene sets representing differentially expressed genes in RNA-seq data from Gar15-13 PDX tumours treated with Decitabine compared to Vehicle (n = 4; FDR < 0.05) **(b)** Volcano plot showing Decitabine *vs*. Vehicle differential expression of genes that are located at enhancer-promoter interactions gained with Decitabine treatment. **(c)** Transcription factor motifs significantly enriched at promoter-interacting enhancers (enhancer OEs) gained with Decitabine treatment. Only motifs with FDR < 0.05 are shown. **(d)** Overlap of consensus ER cistromes in Vehicle and Decitabine-treated Gar15-13 PDX tumours (n = 4 each). Heatmaps indicate ER ChIP-seq signal intensity at ER binding sites gained and lost in Decitabine compared to Vehicle-treated tumours. **(e)** Average signal intensity of ER ChIP-seq binding (Gar15-13 Vehicle and Decitabine tumours) at gained and lost ER binding sites with Decitabine treatment. **(f)** ChromHMM (TAMR) annotation (\**P* value < 0.001) of ER binding sites gained with Decitabine treatment compared to matched random regions across the genome. Size of the overlap is presented in the respective column. **(g)** Transcriptions factor motifs enriched at ER binding sites gained with Decitabine treatment compared to matched random regions generated from ERE binding motifs across the genome. **(h)** Heatmap showing DNA methylation levels (β-values) at gained ER binding sites in Decitabine-treated (n = 4) and Vehicle (n = 4) PDX tumours. **(i)** Browser snapshot of ER ChIP-seq together with EPIC DNA methylation (Vehicle and Decitabine treatment, n = 4 each) showing gain of ER binding and loss of DNA methylation at an enhancer region of ER target gene *ANKRD2*.

To directly address if rewired enhancer-promoter interactions are involved in altered transcription, we identified genes connected to newly gained enhancer OEs and compared their average expression between Vehicle and Decitabine tumours. We identified a total of 4025 genes at new enhancer-promoter interactions (Supplementary Table 6), of which 417 were upregulated after Decitabine treatment (*P* value < 0.05; logFC > 1) (Fig. 4b). Upregulated genes were significantly enriched at gained OE interactions as compared to all genes (Fisher’s exact test *P* value < 2.2e-16) (Fig. 4b). Our data suggests that the dynamic increase in the number of enhancer OEs connected to a promoter results in an overall increase in expression of genes, in agreement with the current models of transcriptional control *via* enhancer-promoter interactions^37,38^.

To further explore the specific role of rewired promoter-anchored interactions in the altered transcriptional program, we evaluated which transcription factors (TFs) are associated with these gained interactions. Notably key transcription factors involved in ER+ breast cancer were highly enriched, including methylation-sensitive ER (estrogen response elements (EREs)) and ELF5 (ETS transcription factor family members), as well as architectural proteins CTCF and ZNF165 (Fig. 4c).

Because of the known role of ER transcription factor in inducing 3D chromatin interactions in ER+ breast cancer cells^14,39–41^ and methylation-sensitive binding^42^, we profiled ER binding site (ERBS) patterns genome-wide in Vehicle (n = 4) and Decitabine-treated (n = 4) tumours via ER ChIP-seq to determine if ER binding was specifically altered by DNA hypomethylation. Differential binding analyses (Diffbind^6^) revealed reprogramming of the ER binding characterised by 1095 gained ERBS and 279 lost ERBS following Decitabine treatment as compared to Vehicle (FDR < 5%) (Fig. 4d) and a stronger average ER ChIP-seq signal at gained ERBS in Decitabine samples as compared to Vehicle, while lost sites showed a moderate decrease in binding intensity genome-wide (Fig. 4e).

Remarkably, over 75% of all gained ERBS were located at distal regulatory regions associated with active and poised enhancers (Fig. 4f and Extended Data Fig. 4b), and these sites were enriched for the estrogen response elements (EREs) DNA motif, followed by FOXA1 (Fig. 4g). Lost ERBS were most frequently positioned close to a TSS (> 40% less than 1Kb from TSS) (Extended Data Fig. 4b), were associated with active promoters (Extended Data Fig. 4c) and were enriched for Sp1 and NFY promoter DNA motifs (Extended Data Fig. 4d). Both lost and gained ERBS were highly enriched for the FOXA1 motif (Fig. 4g and Extended Data Fig. 4d). Additionally, we found a significant loss (approx. 44.4%) of DNA methylation at gained ERBS (Fig. 4h and Extended Data Fig. 4f), as illustrated in Figure 4i at the *ANKRD2* gene locus (Fig. 4i, further example in Extended Data Fig. 4g). In contrast the small proportion of ERBS that were lost remained unmethylated in both Vehicle and Decitabine treated samples (approx. 6.92% DNA methylation change; Extended Data Fig. 4h), suggesting that this subset of ERBS were altered independently of a direct change in DNA methylation.

### Rewired ER-bound chromatin interactions are associated with activation of ER target genes

To determine whether this gain in ER-enhancer binding was associated with rewired 3D chromatin interactions, we integrated the gained ERBS with ectopic enhancer-promoter interactions and associated transcriptional programs. Consistent with ERE motifs enriched at gained enhancer OEs (Fig. 4c), we found significant enrichment for gained ER binding sites (Fig. 5a) and a genome-wide increase in ER binding density at ectopic enhancer OEs induced by Decitabine treatment (Fig. 5b). We propose that these ER-associated enhancer-promoter interactions are mediated by a change in ER binding at enhancer OEs (“ER-bound interactions”).

**Fig. 5.**
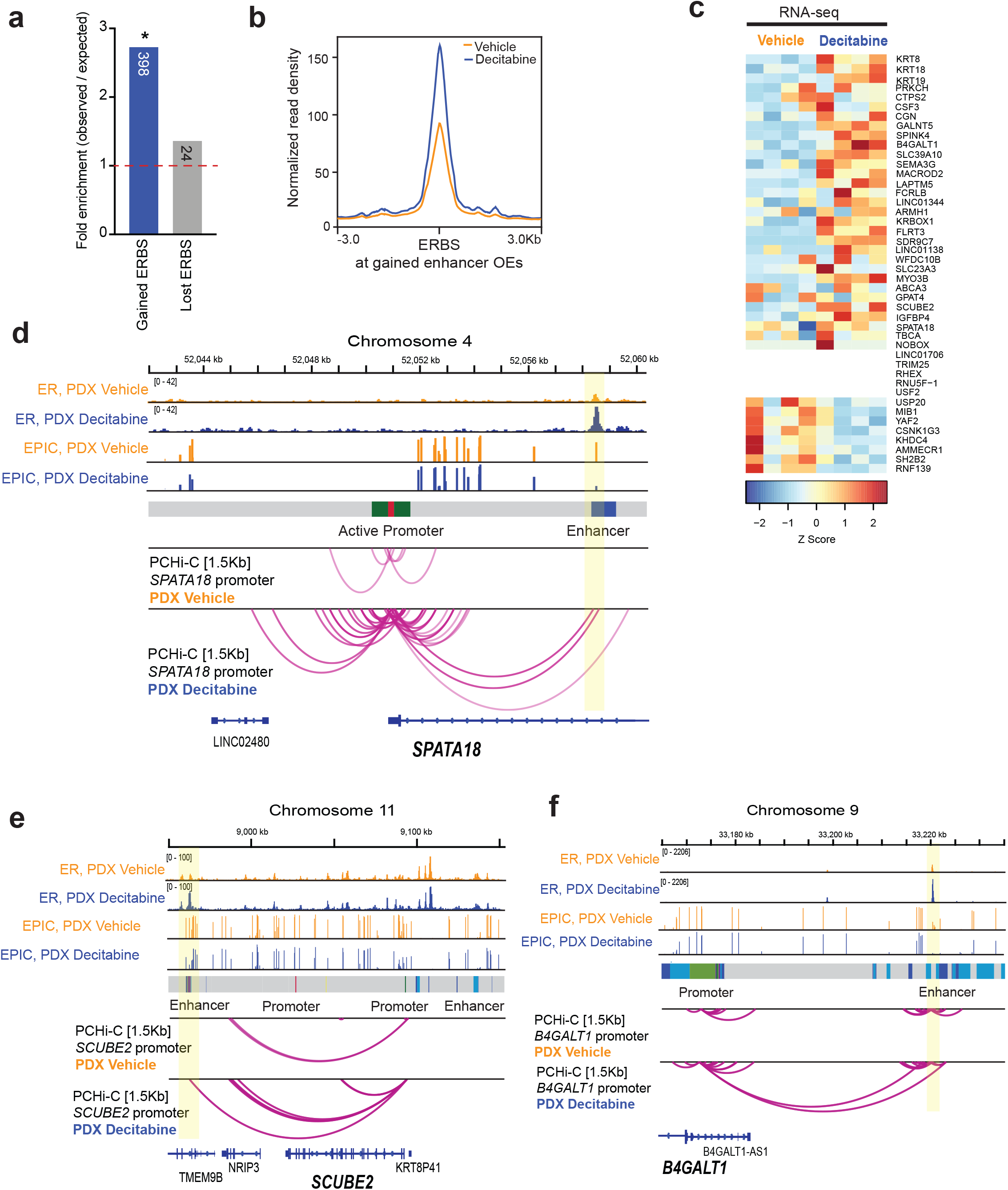
Rewired ER-bound interactions are associated with activation of ER target genes. **(a)** Observed/expected fold change enrichment of gained enhancer OEs for ER binding gained and lost following Decitabine treatment. * *P* value < 0.0001. **(b)** Average ER ChIP-seq signal intensity (Gar15-13 Vehicle and Decitabine-treated tumours) at ER binding sites located at DNA hypomethylation-induced enhancer OEs. **(c)** Heatmap showing expression of genes connected to gained ER-mediated enhancer OEs in Vehicle and Decitabine-treated tumours. **(d)** Browser snapshots showing the promoter-anchored interactions at the *SPATA18* gene, together with the average ER ChIP-seq signal, EPIC DNA methylation and ChromHMM track. Merged replicate data shown (n = 4 each). In Decitabine-treated tumours, the *SPATA18* promoter displays an increased number of interactions with an upstream enhancer region, which gains ER binding with Decitabine treatment, concomitant with loss of DNA methylation. Expression of the *SPATA18* gene was significantly upregulated in Decitabine-treated tumours (shown in Extended Data Fig. 5a). **(e)** Browser snapshots showing promoter-anchored interactions at the *SCUBE2* ER target gene, together with ER ChIP-seq, EPIC DNA methylation and ChromHMM track. Merged replicate data shown (n = 4 each). In Decitabine-treated tumours, the *SCUBE2* promoter displays additional interactions with a distal enhancer, which gains ER binding with Decitabine treatment. Expression of the *SCUBE2* gene was significantly upregulated in Decitabine-treated tumours (shown in Extended Data Fig. 5b). **(f)** Browser snapshots showing promoter-anchored interactions at the *B4GALT1* ER target gene, together with ER ChIP-seq, EPIC DNA methylation and ChromHMM track. Merged replicate data shown (n = 4 each). In Decitabine-treated tumours, the *B4GALT1* promoter displays additional long-range interactions with a distal enhancer, which gains ER binding with Decitabine treatment. Expression of the *B4GALT1* gene was significantly upregulated in Decitabine-treated tumours (shown in Extended Data Fig. 5c).

We next specifically focused on these gained ER-bound enhancer-promoter interactions in detail by identifying connected genes and comparing their expression between Vehicle and Decitabine tumours. The majority (~74%) of these genes showed an overall increase in expression following Decitabine treatment (Fig. 5c) and included established ER target genes (for example *B4GALT1, MYO3B, SEMA3G*) as well as genes associated with a good clinical outcome in ER+ breast cancer (for example *SPATA18, SCUBE2, GALNT5, IGFBP4*). For example, at the *SPATA18* locus, multiple 3D enhancer-promoter interactions are gained with Decitabine treatment, concomitant with gain in ER binding at putative enhancer, loss of DNA methylation and activation of the ER target gene (Fig. 5d and Extended Data Fig. 5a). Moreover, high expression of *SPATA18* gene is associated with good prognosis in ER+ breast cancer (Extended Data Fig. 5a). *SCUBE2* (Fig. 5e and Extended Data Fig. 5b), *B4GALT1* (Fig. 5f and Extended Data Fig. 5c) and *MYO3B* (Extended Data Fig. 5d) genes also exemplify the relationship between Decitabine-induced gain of multiple ER-bound enhancer-promoter interactions and activation of their ER target genes that are associated with good prognosis in ER+ breast cancer.

Taken together, these results reveal a link between Decitabine-induced DNA hypomethylation, rewiring of ER-bound enhancer-promoter interactions and alteration in the ER transcriptional program.

### Dynamics of DNA hypomethylation and re-methylation on 3D chromatin interactions

Finally, to determine the dynamics between DNA methylation alterations and 3D enhancer-promoter rewiring and expression changes, we performed a time-course of Decitabine followed by period of long term recovery in an established cell line model of endocrine-resistance TAMR^14,43,44^ cells. Low dose Decitabine was applied daily for 7 days to induce hypomethylation, followed by a recovery time-point (28 days) to allow for re-methylation of CpG sites demethylated by Decitabine (Fig. 6a). We confirmed loss and recovery of DNMT1 protein expression by Western blot (see Supplementary Note). We assessed changes in DNA methylation, RNA expression and 3D enhancer-promoter interactions on day 7 Decitabine-(“Day 7 Decitabine”) and day 28 post-Decitabine (“Decitabine Recovery”), as well as passage-matched control cells (“Early Control” and “Late Control”) in duplicate. As expected, 7 day Decitabine treatment resulted in widespread DNA hypomethylation in the TAMR cells (approx. 41.84% change in median DNA methylation, *P* value < 0.0001, two-tailed Mann-Whitney test; Fig. 6b and Extended Data Fig. 6a). Substantial genome-wide recovery of DNA methylation was also observed following 28 days of recovery compared to matched vehicle-treated control (approx. 25.46% change in median DNA methylation, *P* value < 0.0001, two-tailed Mann-Whitney test; Fig. 6b and Extended Data Fig. 6a). Similar to the PDX Decitabine-treated samples, we found that DNA hypomethylation changes in TAMR cells, after 7 days of Decitabine treatment, were enriched for ChromHMM ^14^ enhancers (Fig. 6c) and ER binding sites^6^ (Fig. 6d).

**Fig. 6.**
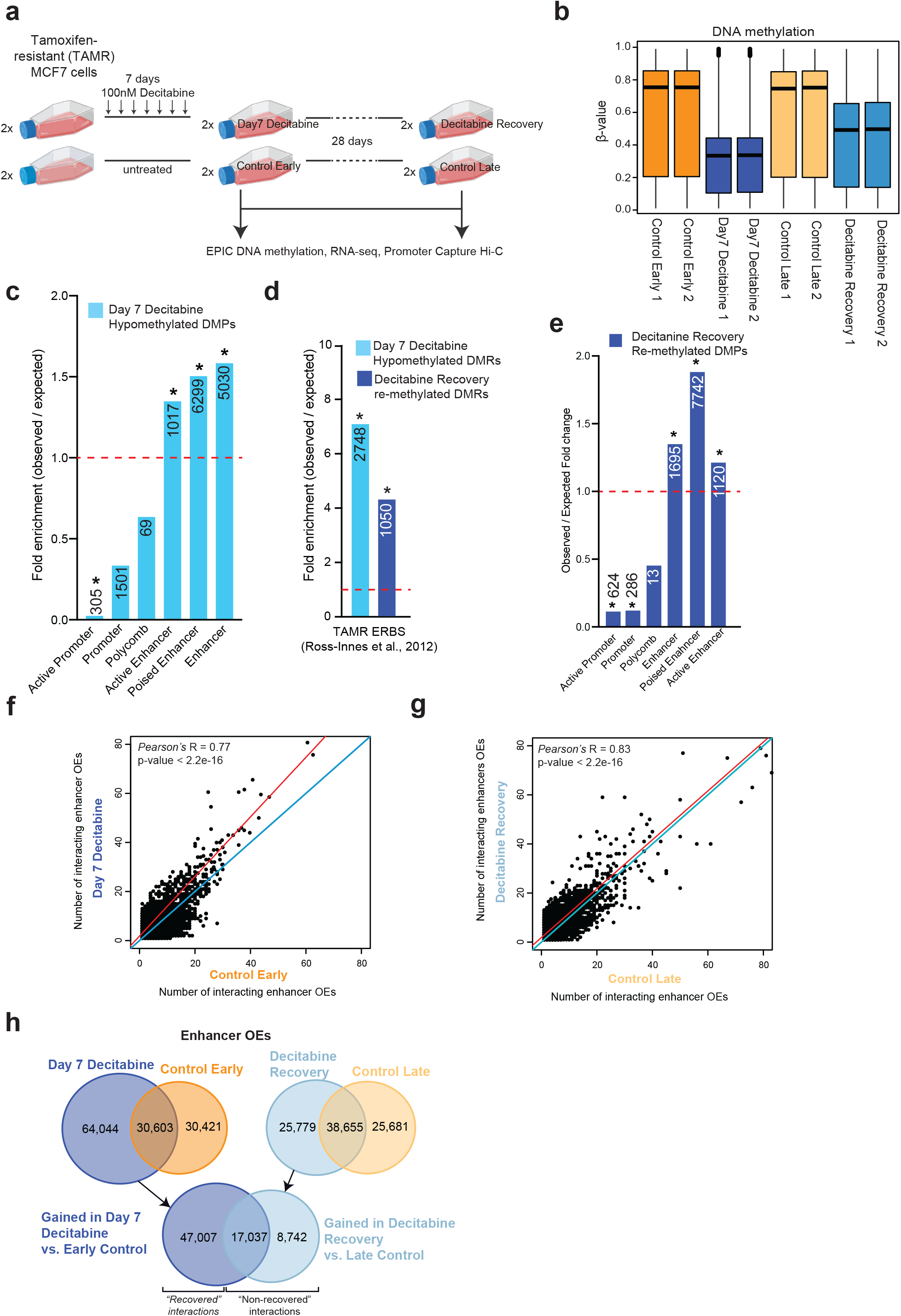
Dynamics between DNA methylation and 3D chromatin interactions. **(a)** Experimental design for the Decitabine treatment loss and DNA methylation “rescue” time-course study in TAMR cell line. **(b)** Boxplots showing the distribution of DNA methylation profiles for Control (Control Early and Control Late; n = 2 replicates), Decitabine-treated (Day 7 Decitabine; n = 2) and 28 days post-Decitabine treatment (Decitabine Recovery; n = 2) TAMR cells for all EPIC probes across the genome. Black line indicates median ± SD. **(c)** Bar plot showing the association of differentially methylated probes in Day 7 Decitabine treatment TAMR cells compared to Control Early TAMR cells across different regulatory regions of the genome as determined by TAMR ChromHMM annotation. Observed over expected fold change enrichment shown. * *P* value < 0.001. The numbers of hypo/hypermethylated probes located within each specific region are presented in the respective column. **(d)** Observed/expected fold change enrichment of Day 7 Decitabine hypomethylated DMRs (as compared to Control Early) and Decitabine Recovery re-methylated DMRs (as compared to Day 7 Decitabine) for ER binding in TAMR cells. * *P* value < 0.0001. **(e)** Bar plot showing the association of EPIC probes that become re-methylated in Decitabine Recovery TAMR cells compared to Day 7 Decitabine cells across different regulatory regions of the genome as determined by TAMR ChromHMM annotation. Observed over expected fold change enrichment shown. * *P* value < 0.001. The numbers of re-methylated probes located within each specific region are presented in the respective column. **(f)** Scatterplot showing number of enhancer OE interactions per promoter bait for each CHiCAGO significant promoter-anchored interaction in Day 7 Decitabine and Control Early TAMR cells. Merged data across replicates shown. **(g)** Scatterplot showing number of enhancer OE interactions per promoter bait for each CHiCAGO significant promoter-anchored interaction in Decitabine Recovery and Control Late TAMR cells. Merged data across replicates shown. **(h)** Overlap between other end (OE) enhancer regions for CHiCAGO significant interactions between Day 7 Decitabine and Control Early samples (left panel) and Decitabine Recovery and Control Late samples (right panel). Below panel shows overlap between gained interactions in Day 7 Decitabine *vs*. Control Early and Decitabine Recovery vs. Control Late, demonstrating the number of “recovered” and “non-recovered” interactions. Merged data across replicates shown.

To study the dynamics of DNA re-methylation on 3D chromatin interactions, we first identified DNA regions that were substantially re-methylated after 28 days of recovery in “Decitabine Recovery” samples, as compared to “Day 7 Decitabine” samples (>30% gain in DNA methylation; Supplementary Table 1). We found that the regions that re-gained DNA methylation were also enriched in poised enhancers and depleted at promoter regions (Fig. 6e). In fact, 10,195 probes located at ChromHMM enhancers hypomethylated after day 7 Decitabine treatment, gained methylation after 28-days recovery (Extended Data Fig. 6b) and were also enriched for ER binding^6^ (Fig. 6d). To determine if 3D enhancer-promoter interactions were also altered in the Decitabine time course, we performed Promoter Capture Hi-C. We found that chromatin interactions separate the control and Decitabine-treated samples on X-axis, with Decitabine Recovery samples clustering together with control samples and away from the Day 7 Decitabine samples on the Y-axis, suggesting substantial “recovery” of chromatin interactions 28 days post Decitabine treatment (Extended Data Fig. 6c). Similar to the PDX results (Fig. 3f), we found that Day 7 Decitabine-treated TAMR samples resulted in a large gain in the number of new enhancer OEs connected to bait promoters (Fig. 6f). Moreover, these additional ectopic interactions were mostly lost in Decitabine Recovery (Fig. 6g). Importantly, the majority of Day 7 Decitabine gained OE enhancers (64,044) interactions were lost in Decitabine Recovery samples (47,007 OE enhancers “recovered” interactions) (Fig. 6h).

We found that gained enhancer-promoter interactions at Day 7 Decitabine treatment, were significantly associated with an overall increase in gene expression (195 up-regulated genes (P value < 0.05; logFC > 1.5; Fisher’s exact test P value < 2.2e-16) (Fig. 7a). No significant increase in expression for genes at Decitabine Recovery interactions was observed (49 up-regulated genes; Fisher’s exact test P value = 0.3167) (Fig. 7b). After 28 days of “recovery” we observed that loss of gene expression was concordant with reversal of ectopic enhancer-promoter interactions, including at key ER-target genes (Fig. 7b). Further evidence of a direct relationship between DNA hypomethylation and ectopic 3D enhancer-promoter interactions is exemplified at ER-target genes also identified in the PDX data: *SPATA18* (Fig. 7c), *B4GALT* (Fig. 7d), *EVL* and *MYO3B* (Extended Data Fig. 6d-e). Notably, we also found a subset of genes that remained upregulated after 28 days of “recovery”, for example *SCUBE2*, where the chromatin contacts were still not fully resolved (Extended Data Fig. 6f).

**Fig. 7.**
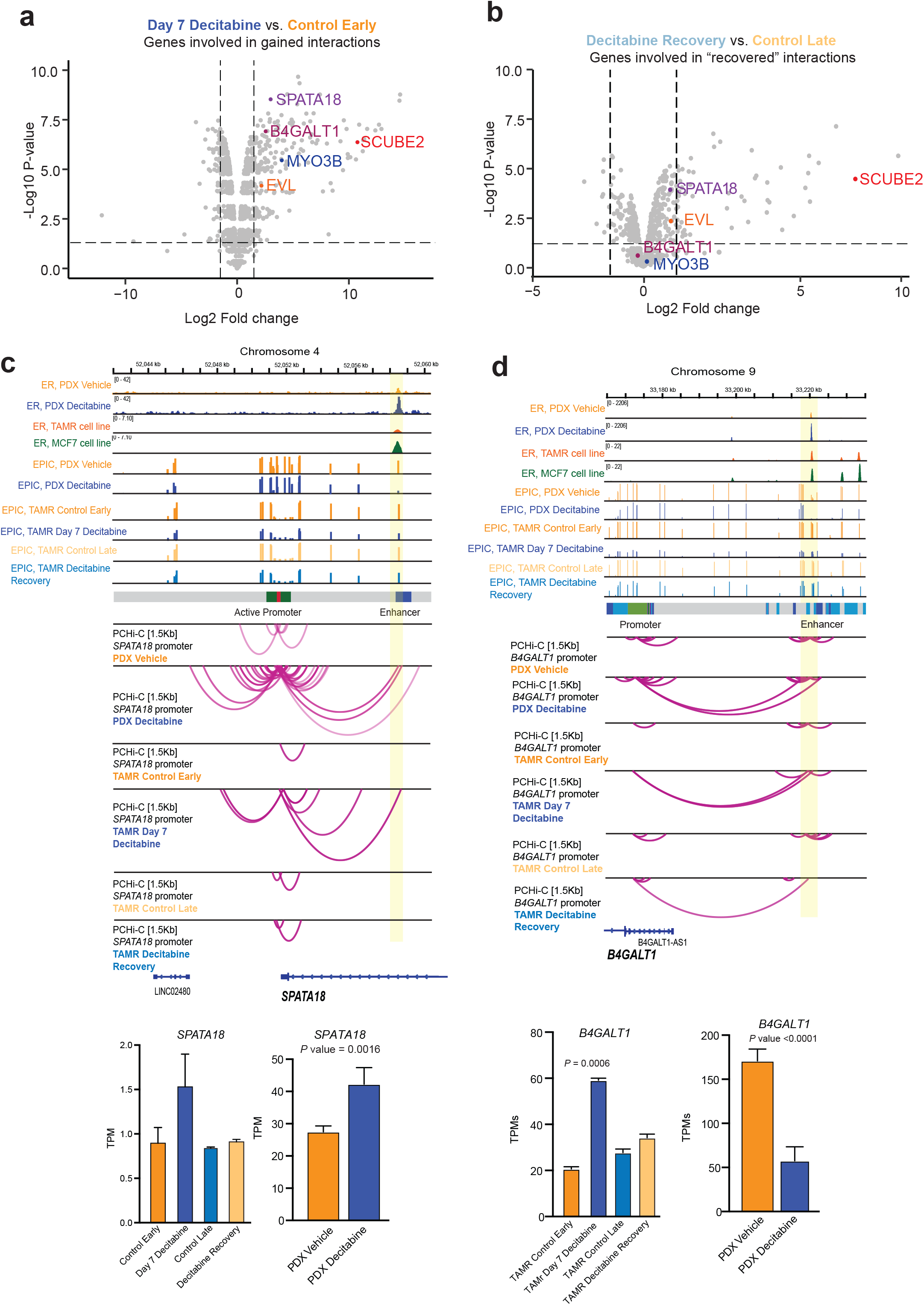
Dynamics of altered ER-bound 3D chromatin interactions on gene transcription. **(a)** Volcano plot showing Day 7 Decitabine *vs*. Control Early differential expression of genes involved in gained differential interactions in Day 7 Decitabine-treated TAMRs. Genes included in representative examples are labelled. **(b)** Volcano plot showing Decitabine Recovery *vs*. Control Late differential expression of genes involved in “recovered” differential interactions in Decitabine Recovery TAMRs. Genes included in representative examples are labelled. **(c)** Browser snapshots showing promoter-anchored interactions at the *SPATA18* ER target gene. Data from Gar15-13 PDX model (ER ChIP-seq, EPIC and PCHi-C) has been overlayed with TAMR cell line data (EPIC and PCHi-C) together with ER ChIP-seq for MCF7/TAMR cell lines (Ross-Innes *et al*., 2012) and ChromHMM track. Merged replicate data shown (n = 4 each for Gar15-13 and n = 2 for TAMRs). In Decitabine-treated PDX tumours and TAMRs (Day 7 Decitabine), the *SPATA18* promoter displays an increased number of interactions with an upstream enhancer region, which gains ER binding with Decitabine treatment in PDXs, concomitant with loss of DNA methylation in both PDXs and TAMRs. These ectopic chromatin interactions are lost after 28 days of recovery with partial recovery of DNA methylation at that locus. Expression of the *SPATA18* gene was significantly upregulated in Day 7 Decitabine-treated TAMRs and suppressed in Decitabine Recovery TAMRs. Expression of the *SPATA18* gene was significantly upregulated in Decitabine-treated *vs*. Vehicle PDXs. **(d)** Browser snapshots showing promoter-anchored interactions at the *B4GALT1* ER target gene. Data from Gar15-13 PDX model (ER ChIP-seq, EPIC and PCHi-C) has been overlayed with TAMR cell line data (EPIC and PCHi-C) together with ER ChIP-seq for MCF7/TAMR cell lines (Ross-Innes *et al*., 2012) and ChromHMM track. Merged replicate data shown (n = 4 each for Gar15-13 and n = 2 for TAMRs). In Decitabine-treated PDX tumours and TAMRs (Day 7 Decitabine), the *B4GALT1* promoter displays an increased number of long-range interactions with a distal enhancer region, which gains ER binding with Decitabine treatment in PDXs, concomitant with loss of DNA methylation in both PDXs and TAMRs. These ectopic chromatin interactions are partially reversed after 28 days of recovery with recovery of DNA methylation at that enhancer locus. *B4GALT1* expression increased in Day 7 Decitabine-treated TAMRs compared to Control Early samples and was restored in Decitabine Recovery TAMRs. Expression of the *B4GALT1* gene was significantly upregulated in Decitabine-treated *vs*. Vehicle PDXs. In TAMRs

## Discussion

Three-dimensional (3D) epigenome remodelling, including widespread changes to DNA methylation and 3D chromatin structure, is an emerging mechanism of gene deregulation in cancer. Our previous work demonstrated that DNA hypermethylation and concomitant loss of ER binding at enhancers was a key event associated with alterations in 3D chromatin interactions in ER+ endocrine-resistant breast cancer^45^. Therefore, we were motivated to determine if these 3D chromatin alterations could be resolved with epigenetic therapies that induce DNA hypomethylation. To date, most studies investigating the use of epigenetic therapies have focused on assessing efficacy in blood cancers and activation of the immune system *via* viral mimicry response. While some studies demonstrated that Decitabine treatment has anti-tumour properties in solid cancers (reviewed in ^46^), the exact mechanism that underpins this response was unclear and evidence for therapeutic activity was inconclusive^21,46^.

Here, using patient-derived xenograft models of ER+ endocrine-resistant breast cancer, we show that treatment with Decitabine induced DNA hypomethylation and had potent anti-tumour activity associated with suppression of tumour growth and cell proliferation gene pathways. To assess the broader functional impact of DNA hypomethylation, we analysed multiple layers of 3D genome organisation, including chromosomal compartments, TADs and 3D chromatin interactions and integrated the 3D data with DNA methylation, transcriptome and ER transcription factor profiles in Decitabine and Vehicle-treated PDX tumours. Collectively our data supports a model (Fig. 8), whereby low dose Decitabine treatment results in enhancer DNA hypomethylation, reprogramming of ER chromatin binding, and rewiring of enhancer-promoter interactions, leading to activation of ER target genes. Importantly, we identified rewired ER-bound chromatin interactions that connect ER-enhancers to specific target genes, which included Estrogen Response Hallmark genes involved in cell cycle inhibition and tumor suppression. Finally, we confirm a mechanistic link between Decitabine-induced DNA hypomethylation, rewiring of 3D chromatin interactions and gene activation using “rescue” DNA methylation experiments in a cell line model of endocrine-resistant breast cancer.

**Fig. 8.**
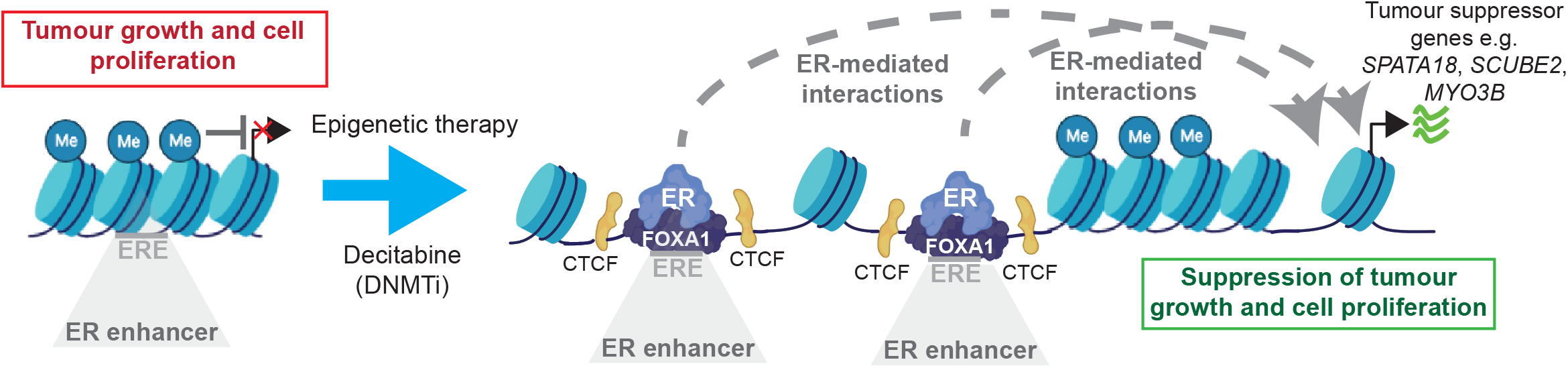
Model of Decitabine-induced 3D epigenome remodelling and suppression of tumour growth. Proposed model of tumour growth suppression induced by epigenetic therapy with Decitabine *via* DNA hypomethylation and subsequent rewiring of ER-bound 3D enhancer-promoter interactions resulting in activation of specific ER target genes.

Decitabine has been previously demonstrated to have some therapeutic efficacy in multiple subtypes of breast cancer and in overcoming drug resistance^4,47^. Transient low-dose treatment with Decitabine resulted in decrease in promoter DNA methylation, gene re-expression, and had antitumor effect *in vivo* in breast cancer cells^48^. Low-dose Decitabine has also been shown to prevent cancer recurrence by disrupting the premetastatic environment through its effect on myeloid-derived suppressor cells in breast and other cancers^49^. In triple negative breast cancer (TNBC) PDX organoids, Decitabine sensitivity was positively correlated with protein levels of DNMTs, suggesting DNMT levels as potential biomarkers of response^50^. A recent study of Decitabine in a panel of breast cancer cell lines observed that Decitabine also induced genes within apoptosis, cell cycle, stress, and immune pathways^51^. However, knockdown of key effectors of the immune pathway did not affect Decitabine sensitivity suggesting that breast cancer growth suppression by Decitabine is independent of viral mimicry^51^.

We found that the low dose Decitabine treatment resulted in minimal DNA hypomethylation at repetitive elements. Despite this, we observed a relatively large number of TEs becoming activated with treatment, consistent with previous studies in cancer cell lines^52^. Loss of DNA methylation at repetitive elements and expression of TEs has been shown to drive viral mimicry response in tumours treated with epigenetic therapies^24–26^. In agreement with studies in colorectal tumour cells^24,53^, our results indicate that treatment with Decitabine results in up-regulation of multiple immune pathways (Interferon and Inflammatory Response pathways), which could promote anti-tumour immunity. However, immunodeficient NOD-*scid IL-2ry^null^* (NSG) mice required for PDX experiments in our study largely lack mature immune cells and therefore the potential immune response could not solely account for the tumour inhibitory effects of Decitabine treatment observed in our study. This highlights the need to study both the immune-based and tumour-based mechanisms that underpin response to epigenetic therapies.

There have been limited studies to date on the role of DNA methylation in shaping the 3D genome organisation^14,16,54–56^. Simultaneous profiling of DNA methylation and 3D genome in single cells revealed pervasive interactions between these two epigenetic layers in regulating gene expression^57^. Additionally, binding of CCCTC-binding factor (CTCF), an insulator protein involved in creating chromatin loops and domain boundaries has been shown to be methylation-sensitive^58–61^. DNA hypermethylation at CTCF binding site in *IDH* mutant gliomas results in its reduced CTCF binding, loss of insulation between topological domains and aberrant gene activation^16^. More recently, DNA hypomethylation has been shown to result in de-compaction of chromatin and loss of compartmental organisation^53,56,62,63^. This is consistent with our data showing closed (B-type) to active (A-type) compartment shifting with Decitabine-induced DNA hypomethylation and reduced interactions between B-type compartments.

A number of studies have suggested an important role of DNA methylation in altering ER binding at regulatory elements^44,64^, however no studies have examined the potential effect on ER-bound 3D chromatin interaction. We have previously shown that DNA methylation differences at enhancers underpin differential ER binding events associated with endocrine resistance^44^. Furthermore, recent studies demonstrated that ER-bound 3D chromatin interactions are altered in endocrine-resistant cells^14,15^. In this study, we show that Decitabine-induced DNA hypomethylation at ER-bound enhancers plays an important functional role in promoting ectopic 3D chromatin interactions and we identify key ER-bound chromatin interactions that control expression of genes associated with endocrine-resistant tumour growth suppression. These results add to the current research, showing that enhancer reprogramming through transcription factor binding or chromatin remodelling can promote transcriptional programs that are associated with endocrine response in breast cancer^65,66^.

Our high-resolution promoter interactions data also revealed an increase in the number of interacting enhancers connecting to gene promoters induced by Decitabine. We hypothesise that the overall increase in enhancer connectivity results in creation of active transcription hubs^38^ or frequently interacting regions^67^ at activated genes, which will be investigated in future studies. This is in agreement with recent reports of transcriptional activation occurring in non-membrane bound nuclear compartments that harbour multiway enhancer-promoter interacting hubs^68^ as well as additive effect of multiway enhancer interactions on gene expression^69^.

In summary, our work highlights a novel molecular mechanism of epigenetic therapies in endocrine-resistant ER+ breast cancer. We provide mechanistic insights into how Decitabine-induced DNA hypomethylation promotes 3D epigenome remodelling including rewiring of ER-mediated 3D enhancer-promoter chromatin interactions. Epigenetic therapy therefore has the potential to overcome cancer therapy resistance by targeting the 3D epigenome architecture to resolve gene deregulation and reduce cancer growth.

## Methods

### Patient-derived xenograft (PDX) models of ER+ breast cancer

All *in vivo* experiments, procedures and endpoints were approved by the Garvan Institute of Medical Research Animal Ethics Committee (HREC #14/35, #15/25) and performed at the Garvan Institute of Medical Research using standard techniques as described previously ^27,70^ in accordance with relevant national and international guidelines. The Gar15-13 model was generated in-house at the St Vincent’s Hospital under the Human Research Ethics protocol (HREC/16/SVH/29) and the HCI-005 model was developed by the Welm laboratory at the Huntsman Cancer Institute (University of Utah)^70^. Gar15-13 was derived from a resected breast cancer liver metastasis of ER-positive, PR-negative, HER2-negative metastatic breast cancer^28^. HCI-005 was derived from a pleural effusion of ER-positive (ERmutL536P), PR-positive, HER2-positive metastatic breast cancer. Growth of HCI-005 was supported by estrogen supplementation in the form of a 60 day 17-β-estradiol pellet implanted simultaneously with the tumour chunks. Mice implanted with Gar15-13 did not receive estrogen supplementation as this model does not require additional estrogen for growth^28^.

At surgery, 4 mm^3^ sections of tumour tissue were implanted into the 4th inguinal mammary gland of 6–8-week-old female NOD-*scid* IL2Rγ^null^ (NSG) mice, obtained from Australian BioResources (Sydney, Australia). For HCI-005, tumour growth was supported by implantation of an E2 pellet inserted subcutaneously via the incision site before it was sealed with an Autoclip wound clip. When tumours became palpable, tumour growth was assessed twice weekly by calliper measurement (using the formula: width^2^ × length/2) and mice were randomised to treatment arms when tumours reached 200 mm^3^ using an online randomisation tool (https://www.graphpad.com/quickcalcs/randomize1.cfm) (n = 6 - 8 mice per group for therapeutic studies, exact numbers specified in figure legends).

### Pharmacological treatments in PDX models

The DNA methyltransferase inhibitor, Decitabine (5-Aza-2’-deoxycytidine: Sigma, Cat #3656) was reconstituted in PBS and stored at −80°C.

Decitabine was administrated in 0.5mg/kg/mouse dose in PBS 100ul IP 2 times weekly. Vehicle mice were treated with 100μl PBS IP. Mice were treated for 60 days or until tumour volume reached 1000 mm^3^. Upon reaching ethical or pre-defined experimental endpoints, mice were euthanized, and the primary tumour collected. After weighing, the tumour was cut into pieces that were allocated to be snap-frozen, fixed overnight at 4°C in 10% neutral-buffered formalin, or embedded in cryo-protective optimal cutting temperature (OCT) compound before being snap frozen. Frozen samples were kept at −80°C. Formalin-fixed samples were sent to the Garvan Institute Histology Core Facility for paraffin embedding. Tumour growth curves were analysed in GraphPad Prism (GraphPad Software) by two-tailed, unpaired t-test. Tumour mass at end-point was analysed by two-tailed Mann-Whitney t-test as per figure legends unless otherwise specified.

### Cell culture

MCF7 breast cancer cells and the corresponding endocrine-resistant sub-cell lines were kindly given to our laboratory by Dr. Julia Gee (Cardiff University, UK). Tamoxifen-resistant MCF7 cells (TAMR) were previously generated by the long-term culture of MCF7 cells in phenol red-free RPMI medium containing 10% charcoal-stripped FBS (Gibco) and 4-OH-tamoxifen (1 × 10^-7^ M; TAM).

### Pharmacological treatments in cell lines

Cells were treated daily with Decitabine (100nM) for 7 consecutive days. After 7 days, fresh media was added, and cells were harvested at day 7 (“Decitabine Day 7”). Control cells were cultured for a total of 11 days in normal media and harvested for “Control Early” on day 11. For the “Decitabine Recovery” samples, cells were treated daily with Decitabine (100nM) for 7 consecutive days, after which fresh media was added; cells were cultured for 21 additional days and harvested on day 28 (“recovery” - reintroduction of DNA methylation). Matched control cells were cultured for 28 days in normal media and harvested at day 28 for “Control Late”. DNMT1 protein levels were confirmed by Western blot (see Supplementary Note).

### Cellular viability assay

Tamoxifen-resistant (TAMR) MCF7 cells were seeded in a white flat-bottom 96-well plate at a density of 2500 cells/well based on doubling time and allowed to adhere overnight. Cells were treated daily with Decitabine (0nM, 10nM, 50nM, 100nM, 200nM, 300nM, 500nM, 1uM, 5uM, 10uM and 100uM) for 72h and cellular viability was assessed on day 5, following 2 days of recovery using the alamarBlue (ThermoFisher, DAL1025) assay in accordance with the manufacture’s recommendations (see Supplementary Note).

### Immunohistochemistry (IHC) and quantification

Tumour tissue was harvested and immediately fixed in 10% neutral buffered formalin at 4°C overnight before dehydration and paraffin embedding. Antibodies used for IHC were anti-ER (M7047, 1:300, Agilent) and anti-Ki67 (M7240, 1:400, Agilent). Primary antibodies were detected using biotinylated IgG secondary antibodies (Agilent, 1:400), using streptavidin-HRP (Agilent) for amplification of signal followed by the addition of 3,3’-diaminobenzidine (Sigma) substrate. Images were scanned using Leica Aperio Slide Scanner (Leica Biosystems) and analysed using QuPath software to differentiate tumour tissue from stroma and necrosis, and to quantify Ki-67 positivity in tumour tissue.

### Low input *in situ* Hi-C in snap-frozen PDX tumour samples and TAMR cells

Tumour tissue samples were flash frozen and pulverized in liquid nitrogen before formaldehyde cross-linking in TC buffer. Hi-C was then conducted using the Arima-HiC kit, according to the manufacturer’s protocols (Cat. #A510008) with minor modifications. Briefly, for each Hi-C reaction between ~100,000 - 500,000 cells were cross-linked with 2% formaldehyde and nuclei were isolated by incubating cross-linked cells in Lysis Buffer at 4°C for 30 minutes. The Arima kit uses two restriction enzymes recognizing the following sequence motifs ^GATC and G^ANTC (N can be either of the 4 genomic bases), which after ligation of DNA ends generates 4 possible ligation junctions in the chimeric reads: GATC-GATC, GANT-GATC, GANT-ANTC, GATC-ANTC. Hi-C libraries were prepared using the Swift Biosciences Accel-NGS 2S Plus DNA Library Kit with a modified protocol provided by Arima with 8 PCR cycles for library amplification as required. Hi-C libraries were sequenced on Illumina HiSeq X10 in 150bp paired-end mode.

### Promoter Capture Hi-C

To perform promoter Capture Hi-C (PCHi-C), we computationally designed RNA probes that capture promoter regions of previously annotated human protein-coding genes. Promoter Capture was performed as previously described^71–73^ using the Arima HiC+ kit for Promoter CHi-C (human) (Cat #A510008, #A303010, #A302010 and #A301010). First, to identify promoter capture targets, 23,711 unique Ensembl annotated genes were extracted from the GRCh38 gene annotation file in Ensembl database, version 95. These compromised of protein-coding (18,741), antisense (84), lincRNA (170), miRNA (1,878), snoRNA (938), snRNA (1,898) or multiple (2) transcripts. We then located transcription start sites (TSS) of each gene and mapped the TSS coordinates to the in silico digested genome (^GATC and G^ANTC) and extracted the restriction fragment containing the TSS, as well as one restriction fragment upstream and one restriction fragment downstream for each TSS. The final target list of TSS mapped to 3 consecutive restriction fragments. The average length of the 3 consecutive restriction fragments for each TSS is 786bp and the median is 927bp, with a range of 54-4174bp.

Moreover, for the individual restriction fragments smaller than 700bp, all nucleotides within these fragments are less than or equal to 350bp from the nearest cut site, and therefore the entire restriction fragment was defined as a target region for subsequent probe design. This scenario represents the vast majority of cases, because the mean length of an individual restriction fragment is 263bp, with a median of 431bp. However, if an individual restriction fragment was greater than 700bp, then the 350bp on each inward facing edge of the restriction fragment was defined as a target region for probe design, and the centre most portion of the restriction fragment was excluded from the probe design. After this final processing, a final BED file of target bait regions was input into the Agilent SureDesign tool, and probes were designed using a 1X tiling approach, with moderate repeat masking and balanced boosting. Promoter Capture was carried out using Hi-C libraries from three Vehicle-treated tumour samples and three Decitabine-treated tumour samples with the SureSelect target enrichment system and RNA bait library according to manufacturer’s instructions (Agilent Technologies kit), using 12 post-capture PCR cycles as required. PCHi-C libraries were sequenced on the Illumina HiSeq X10 platform in 150bp paired-end mode.

### Microarray genome-wide DNA methylation

DNA from four Decitabine and four Vehicle-treated tumours from two PDX models (Gar15-13 and HCI-005) was isolated from snap-frozen tumour samples using the Qiagen QIAamp DNA Mini Kit. DNA (500ng) was treated with sodium bisulphite using EZ-96 DNA methylation kit (Zymo Research CA, USA). DNA methylation was quantified using the Illumina InfiniumMethylationEPIC (EPIC) BeadChip (Illumina, CA, USA) run on the HiScan System (Illumina, CA, USA) using manufacturer’s standard protocol.

### ChIP-seq

Tumour samples were snap-frozen in Optimal Cutting Temperature compound (Tissue-Tek) and used for ER ChIP-seq experiments. Using a cryostat (Leica, #CM3050-S), a minimum of 50 x 30 μm sections were cut from each tumour at −20 °C and subjected to double cross-linking with DSG and formaldehyde as previously described^30^. ER ChIP-seq was performed with an anti-ER antibody (Santa Cruz, SC-543X). 5μg of antibody was used to ChIP each tumour sample and 10ng of immunoprecipitated DNA was submitted to the David R. Gunn Genomics Facility at the South Australian Health and Medical Research Institute (SAHMRI) for sequencing. Conversion of the DNA into sequencing libraries was performed using the Ultralow Input Library Kit (Qiagen, #180495) and sequenced on the Illumina NextSeq 500 (Illumina) in 75bp single-end mode to achieve a minimum of 20 million reads per sample.

### RNA-seq

RNA was extracted from snap-frozen tumour PDX tissue and TAMR cell line samples using the RNeasy Mini Kit (QIAGEN) and quality of purified RNA was confirmed with RNA ScreenTape TapeStation (Agilent). All samples processed for RNA-seq had a RIN equivalent (RIN^e^) quality score ≥ 8.0. Total RNA was supplied to the genomics core facility (Kinghorn Centre for Clinical Genomics) for library preparation and sequencing. RNA was prepared for sequencing using the TruSeq Stranded mRNA Library Prep kit (Illumina) and libraries were sequenced on Illumina NovaSeq 6000 S4 in paired-end mode.

### EPIC DNA methylation analyses

Raw intensity data (IDAT) files were imported and quality controlled using *minfi* package (v.1.34.0)^74^. Data was then normalised with background correction. To reduce the risk of false discoveries, we removed probes affected for cross-hybridization to multiple locations on the genome or overlapped SNPs, as previously described^75^. Beta (β) values were calculated from unmethylated (U) and methylated (M) signal [M/(U + M + 100)] and ranged from 0 to 1 (0 to 100% methylation). β values of loci whose detection P values were > 0.01 were assigned NA in the output file. To map EPIC arrays to hg38/GRCh38 assembly, all probes were annotated with the EPIC.hg38.manifest.tsv.gz files as described in^76^.

For initial visualisation of the EPIC data, multidimensional scaling plots were generated using the ‘mdsPlot’ function in the *minfi* Bioconductor package (v.1.34.0)^74^. Differential analyses were then performed between treatment arm with Decitabine versus Vehicle samples. For each comparison, β values were transformed using logit transformation: *M* = log2(β/(1-β)). We used the limma package (v.3.46)^77^ to identify DMPs between sample groups with adjusted *P* value cut-off of < 0.01. The R package DMRcate (v.2.2.3)^78^ was used to identify DMRs, with DMP *P* value cut-offs of FDR < 0.01. DMRs were defined as regions with a maximum of 1000 nucleotides between consecutive probes and a minimum of 2 CpG sites, a methylation change > 30% and we applied Benjamini-Hochberg correction for multiple testing. ChromHMM data downloaded from GEO (GSE73783) for tamoxifen-resistant (TAMR) MCF7 cells was used to annotate DMPs to chromatin states. REMP R package (v.1.14.0)^32^ was used to assess genome-wide locus-specific DNA methylation of repeat elements (LTR, LINE1 and Alu) from EPIC data with IlluminaHumanMethylationEPICanno.ilm10b5.hg38 annotation (GitHub).

### Hi-C analyses

Hi-C sequenced reads (150bp paired-end) were quality checked with FastQ Screen v.0.14.1^79^ for mouse host reads contamination. Reads were then processed with Xenome (v.1.0.1)^80^ as described in ^81^. Remaining reads were aligned to the human genome (hg38/GRCh38) using HiC-Pro^82^ (v.2.11.4). Initially, to generate Hi-C contact matrices, the aligned Hi-C reads were filtered and corrected using the ICE “correction” algorithm^83^ built into HiC-Pro, which corrects for the CNV-related variability in the tumours^84^. Inter-chromosomal interactions were excluded from further analyses to control for the effect of inter-chromosomal translocations in the tumours. Contact matrices for 3D genome feature annotation and visualisation were created and Knight-Ruiz (KR) normalized using Juicer tools^85^ using contact matrices in .hic format generated by hicpro2juciebox script in HiC-Pro as input (hic file version 8). Hi-C matrices were visualised using JuiceBox^86^ and WashU Epigenome Browser^87^. We confirmed data quality by assessing the proportion of cis/trans interactions and percentage of valid fragments for each library. Overall, we obtained an average of 100 million unique, valid contacts per replicate (~310 million per treatment arm), for an average resolution of 10kb. Statistics for each library can be found in Supplemental Table 2. These data was used to derive loops, TAD boundaries and chromosomal compartment structures.

### Insulation score and identification of TAD boundaries

Topological domain boundary calling was performed by calculating insulation scores in ICE normalised contact matrices at 20kb resolution using TADtool^88^. To identify appropriate parameters, we called TADs across chromosome 1 using contact matrices at 20kb and threshold values: 10, 50 and 100. The final TADs were called for all chromosomes at window 102353 and cut-off value 50. Boundaries that were found overlapping by at least 1 genomic bin were merged. Boundaries separated by at least one genomic bin were considered different between datasets. Pyramid-like heatmap plots were generated with GENOVA^89^.

### Identification of compartments A and B

For each chromosome in each sample, compartments where called using the standard PCA method^33^ in Homer package (version 4.8). The resolution was set to 50kb and the window size to 100 kb. Compartments were defined as regions of continuous positive or negative PC1 values using the findHiCCompartments.pl tool in Homer. To detect which compartment is the open “A-type” and which is the closed “B-type,” the genome-wide gene density was calculated to assign the “A-type” and “B-type” compartmentalisation. To identify genomic regions that switch between two compartment types, we used correlation difference script (getHiCcorrDiff.pl) with findHiCCompartments.pl tool in Homer. Compartments were considered common if they had the same compartment definition within the same genomic bin. Compartment changes between treatment arms were computed after considering compartments that were overlapping between biological replicates, unless otherwise indicated.

To directly quantify the tendency of each region to interact with the other regions in either A or B compartments, we calculated the “A/B interaction ratio”, defined for each 100kb genomic window as the ratio of interaction frequency with A versus B compartments using O/E matrix with GENOVA^89^ (v1.0.0, https://github.com/robinweide/GENOVA). Log2 contact enrichments were plotted as a heat saddle plot. Summarised A-A, B-B and B-A compartment strengths were calculated as the mean log2 contact enrichment between the top (A) or bottom (B) 20% of PC1 percentiles. The compartment strength ratio was calculated as log2(A-A/B-B).

### PCHi-C analyses

PCHi-C sequenced reads were mapped and filtered using HiCUP (v.0.7.4)^90^ with hg38/GRCh38 genome digested with –arima flag and minimum di-tag length set to 20. Statistics for each library can be found in Supplementary Table 3. On target rate was calculated by counting number of valid, unique reads overlapping bait fragments (min. overlap > 0.6). Unique, valid mapped reads from HiCUP were converted into .chinput files using bam2chicago.sh utility and obtained chinput files were further filtered and processed with CHiCAGO (v.1.14.0)^34^. CHiCAGO design files were created with following parameters to account for multiple restriction enzymes used in the Arima HiC kit and the Arima-specific design of the bait fragments: MaxLBrowndist = 75000; binsize = 1500; minFragLen = 25; maxFragLen = 1200. Significant interactions were called with CHiCAGO using score cut-off of 5. All bait-to-bait interactions were discarded. Chicdiff package^35^ (v.0.6) was used to compare PCHi-C data from Vehicle and Decitabine tumours and the difference in the mean asinh-transformed CHiCAGO scores between conditions above 1 was used to prioritise the potential differential promoter-anchored interactions. Only interactions that have CHiCAGO score of more than 5 in at least 2 replicates were included for downstream analysis. Volcano plots were generated using EnhancedVolcano R package (v.1.8.0)^91^. For downstream analysis of merged replicate data and for visualisation of interactions in WashU Epigenome Browser^87^ replicates were merged with CHiCAGO. We defined reprogrammed enhancer-promoter interactions by constructing a consensus, gained and lost subset of promoter-anchors (baits) and other end anchors (OEs) based on CHiCAGO promoter interactions, Chicdiff analysis and setdiff R function across the replicates. The following criteria was used to obtain these regions: CHiCAGO score > 5 in 2 out of 3 replicates in either condition, Chicdiff generated asinh-transformed CHiCAGO scores between conditions above 1 and no overlap between regions, allowing for 10Kb maximal gap in 3 out of 3 replicates. ChromHMM data downloaded from GEO (GSE73783) for tamoxifen-resistant (TAMR) MCF7 cells was used to annotate promoter-anchored interactions to chromatin states.

### RNA-seq data analyses

For canonical gene expression, RNA-seq raw reads were quality checked (FastQ Screen v.0.14.1^79^). Sequence adaptors were trimmed using Trim Galore (v.0.11.2), reads were processed with Xenome v.10.1.^80^ to remove mouse sequences and remaining reads were mapped with STAR (v.2.7.7a)^92^ to the hg38/GRCh38 human genome build with GENCODE v33 used as a reference transcriptome (parameter settings: −quantMode TranscriptomeSAM-outFilterMatchNmin 101 −outFilterMultimapNmax 20). Statistics for each library can be found in Supplementary Table 4. TMM normalisation was applied to normalise for RNA composition^93^ and differential expression was performed with edgeR 3.18.1 ^94^ using the generalised linear model (GLM). RNA-seq tracks were generated using bedtools v.2.22 genomeCoverageBed to create normalised.bedGraph files and bedGraphToBigWig (USCS utils) to create.bigwig files.

For analysis of TE expression, adapter-trimmed, human-only RNA-seq reads were mapped with STAR (v.2.7.7a), allowing for multimapping alignments (flags: -- outFilterMultimapNmax 100 --outFilterMismatchNmax 100). Annotation GTF files for canonical genes were downloaded from Ensembl genome browser (v.102, GRCh38.p13 assembly) and TE annotation GTF file (GRCm38_Ensembl_rmsk_TE.gtf) was downloaded from TEtranscripts ^95^ website (http://hammelllab.labsites.cshl.edu/software/#TEtranscripts). Normalisation and differential expression was performed for all genes and TEs using DESeq2^96^. TEs with and FDR < 0.1 and logFC > 0.7 were considered differentially expressed in pair-wise comparisons. Volcano plots were generated using EnhancedVolcano R package (v.1.8.0)^91^.

### ChIP-seq data analyses

ChIP-seq reads were aligned against human genome (hg38/GRCh38) using bowtie2 with default parameters^97^. Non-uniquely mapped, low quality (MAPQ < 15) and PCR duplicate reads were removed. Peak calling of individual ChIP–seq experiments was performed with MACS2 with default parameters^98,99^. Statistics for each library can be found in Supplementary Table 5. Consensus peaks were identified by intersecting MACS2 peaks obtained from each sample using bedtools intersect (v.2.25.0) with min. overlap > 0.6. Differential binding analyses were performed using DiffBind (v.3.0.9)^6^ and DESeq2 (v.1.3.0)^96^ with FDR < 5%. Enrichment analyses were performed using GAT^100^, ChIPseeker (v.1.26.0)^101^ and normalised to library size. Merged bigwig tracks for visualisation were created from merged bam files from all replicates using the bamCoverage function with scaling factor normalisation and heatmaps and average profiles were plotted with deepTools2^102^. ChromHMM data downloaded from GEO (GSE73783) for tamoxifen-resistant (TAMR) MCF7 cells was used to annotate ER binding sites to chromatin states. The Homer motif discovery suite^103^ was used for motif analysis using the bed regions from differential peaks detected by DiffBind with default parameters, using random, matched regions as background. Merged bigwig tracks for visualisation were created from merged bam files from all replicates using the bamCoverage function with scaling factor normalisation and heatmaps and average profiles were plotted with deepTools2^102^.

### Gene ontology analysis

Gene ontology enrichment analysis and pathway enrichment were done using GSEA (v.4.1.0) and MSigDB 7.2^36^. All significant biological processes and pathways had an adj. P value < 0.001.

### Statistical analyses

The Mann-Whitney-Wilcoxon test was used for 2-group non-parametric comparisons. Unless otherwise stated, statistical tests were two-sided. Permutation test was used to calculate empirical *P* values, which does not make any assumptions on the underlying distribution of the data. Tumor growth curve data was analysed at ethical end point using a two-tailed unpaired Student’s *t*-test. IHC data were analysed by a two-tailed, unpaired Student’s *t*-test.

### Public datasets

ChIP-seq data sets were downloaded from GSE32222 by Ross-Innes *et al*., 2012^6^. ChromHMM data was downloaded from GSE73783 by Achinger-Kawecka *et al*., 2020^14^.

## Supporting information

Supplementary Notre

## Data availability

All sequencing data created in this study have been uploaded to the Gene Expression Omnibus (GEO; https://www.ncbi.nlm.nih.gov/geo/) and are available under primary accession code GSE171074 and GSE216989. Biological material used in this study can be obtained from the authors upon request.

## Code availability

Python script language (v.2.7.8 and v.3.9.1) and R (v.3.6.3 and v.4.0.3) were used to develop the bioinformatics methods and algorithms in this work. All code for Hi-C and PCHi-C analyses is available within the GitHub repository https://github.com/JoannaAch/PDX_Decitabine_3DEpigenome.

## Acknowledgments

We thank members of the Clark Laboratory for helpful discussions and careful reading of the manuscript. We thank Arima Genomics Inc. for advice with performing Hi-C and Promoter Capture Hi-C in snap-frozen tumour tissues. S.J.C. is a National Health and Medical Research Council (NHMRC) Senior Principal Research Fellow #1063559. J.A-K. is a National Breast Cancer Foundation (NBCF) Mavis Robertson Fellow #IIRS-21-047. E.L. is a NBCF Endowed Chair. T.E.H. is an NBCF Fellow #IIRS-19-009. C.S. is an NBCF Fellow IIRS-18-137. Q.D. is a NHMRC Investigator Grant recipient #1177792. J.M.W.G is supported by a Breast Cancer Now Fellowship and the Tenovus Cancer Charity. This research was supported by the NHMRC Project Grant #1128916 (to S.J.C), Movember & National Breast Cancer Foundation Collaboration Initiative grant (MNBCF-17-012 to T.E.H., E.L., S.J.C.) and the Cancer Council NSW grants RG16-09 (to S.J.C) and RG20-04 (to J.A-K.). Research funding (to S.J.C.) was provided by Van Andel Institute through the Van Andel Institute – Stand Up To Cancer Epigenetics Dream Team. Stand Up To Cancer is a division of the Entertainment Industry Foundation, administered by AACR. The contents of the published material are solely the responsibility of the administering institution and individual authors and do not reflect the views of the NHMRC.

## Author Contributions

Conception (S.J.C., C.S., J.A-K) and Experimental Design: J.A-K., K-M.C., N.P., T.E.H., E.L., C.S. and S.J.C. Methodology and Data Acquisition: J.A-K., K-M.C., N.P., E.C., A.Y., A.W., G.L.L., A.S., E.W., J.G., A.K., S.A., J.S. Data Analysis: J.A-K., E.C., H.H.M., B.M., S.C. and Q.D. Data Interpretation: J.A-K., K-M.C., N.P., E.C., E.C.C., Q.D., T.E.H., E.L., C.S. and S.J.C. Funding Acquisition: J.A-K., T.E.H., E.L. and S.J.C. Manuscript Writing: J.A-K. and S.J.C. with input from all authors.

## Ethics declarations

A.S. is an employee of Arima Genomics, Inc. Other authors declare no competing interests.

## References

1. Farcas, A.M., Nagarajan, S., Cosulich, S. & Carroll, J.S. Genome-Wide Estrogen Receptor Activity in Breast Cancer. Endocrinology 162 (2021).

2. Musgrove, E.A. & Sutherland, R.L. Biological determinants of endocrine resistance in breast cancer. Nat Rev Cancer 9, 631–43 (2009).

3. Ali, S. & Coombes, R.C. Endocrine-responsive breast cancer and strategies for combating resistance. Nat Rev Cancer 2, 101–12 (2002).

4. Garcia-Martinez, L., Zhang, Y., Nakata, Y., Chan, H.L. & Morey, L. Epigenetic mechanisms in breast cancer therapy and resistance. Nat Commun 12, 1786 (2021).

5. Dimitrakopoulos, F.I., Kottorou, A. & Tzezou, A. Endocrine resistance and epigenetic reprogramming in estrogen receptor positive breast cancer. Cancer Lett 517, 55–65 (2021).

6. Ross-Innes, C.S. et al. Differential oestrogen receptor binding is associated with clinical outcome in breast cancer. Nature 481, 389–93 (2012).

7. Lupien, M. et al. Growth factor stimulation induces a distinct ER(alpha) cistrome underlying breast cancer endocrine resistance. Genes Dev 24, 2219–27 (2010).

8. Achinger-Kawecka, J. & Clark, S.J. Disruption of the 3D cancer genome blueprint. Epigenomics 9, 47–55 (2017).

9. Taberlay, P.C. et al. Three-dimensional disorganization of the cancer genome occurs coincident with long-range genetic and epigenetic alterations. Genome Res 26, 719–31 (2016).

10. Rhie, S.K. et al. A high-resolution 3D epigenomic map reveals insights into the creation of the prostate cancer transcriptome. Nat Commun 10, 4154 (2019).

11. Barutcu, A.R. et al. RUNX1 contributes to higher-order chromatin organization and gene regulation in breast cancer cells. Biochim Biophys Acta 1859, 1389–1397 (2016).

12. Barutcu, A.R. et al. SMARCA4 regulates gene expression and higher-order chromatin structure in proliferating mammary epithelial cells. Genome Res 26, 1188–201 (2016).

13. Barutcu, A.R. et al. Chromatin interaction analysis reveals changes in small chromosome and telomere clustering between epithelial and breast cancer cells. Genome Biol 16, 214 (2015).

14. Achinger-Kawecka, J. et al. Epigenetic reprogramming at estrogen-receptor binding sites alters 3D chromatin landscape in endocrine-resistant breast cancer. Nat Commun 11, 320 (2020).

15. Zhou, Y. et al. Temporal dynamic reorganization of 3D chromatin architecture in hormone-induced breast cancer and endocrine resistance. Nat Commun 10, 1522 (2019).

16. Flavahan, W.A. et al. Insulator dysfunction and oncogene activation in IDH mutant gliomas. Nature 529, 110–4 (2016).

17. Kloetgen, A. et al. Three-dimensional chromatin landscapes in T cell acute lymphoblastic leukemia. Nat Genet 52, 388–400 (2020).

18. Vilarrasa-Blasi, R. et al. Dynamics of genome architecture and chromatin function during human B cell differentiation and neoplastic transformation. Nat Commun 12, 651 (2021).

19. Corces, M.R. & Corces, V.G. The three-dimensional cancer genome. Curr Opin Genet Dev 36, 1–7 (2016).

20. Yang, Y. et al. The 3D Genomic Landscape of Differential Response to EGFR/HER2 Inhibition in Endocrine-Resistant Breast Cancer Cells. Biochim Biophys Acta Gene Regul Mech, 194631 (2020).

21. Bates, S.E. Epigenetic Therapies for Cancer. N Engl J Med 383, 650–663 (2020).

22. Pandiyan, K. et al. Functional DNA demethylation is accompanied by chromatin accessibility. Nucleic Acids Res 41, 3973–85 (2013).

23. Yang, X. et al. Gene body methylation can alter gene expression and is a therapeutic target in cancer. Cancer Cell 26, 577–90 (2014).

24. Mehdipour, P. et al. Epigenetic therapy induces transcription of inverted SINEs and ADAR1 dependency. Nature 588, 169-+ (2020).

25. Roulois, D. et al. DNA-Demethylating Agents Target Colorectal Cancer Cells by Inducing Viral Mimicry by Endogenous Transcripts. Cell 162, 961–973 (2015).

26. Loo Yau, H. et al. DNA hypomethylating agents increase activation and cytolytic activity of CD8(+) T cells. Mol Cell (2021).

27. Cazet, A.S. et al. Targeting stromal remodeling and cancer stem cell plasticity overcomes chemoresistance in triple negative breast cancer. Nat Commun 9, 2897 (2018).

28. Chia, K. et al. Non-canonical AR activity facilitates endocrine resistance in breast cancer. Endocr Relat Cancer 26, 251–264 (2019).

29. Portman, N. et al. MDM2 inhibition in combination with endocrine therapy and CDK4/6 inhibition for the treatment of ER-positive breast cancer. Breast Cancer Research 22(2020).

30. Hickey, T.E. et al. The androgen receptor is a tumor suppressor in estrogen receptor-positive breast cancer. Nat Med 27, 310–320 (2021).

31. DeRose, Y.S. et al. Tumor grafts derived from women with breast cancer authentically reflect tumor pathology, growth, metastasis and disease outcomes. Nat Med 17, 1514–20 (2011).

32. Zheng, Y. et al. Prediction of genome-wide DNA methylation in repetitive elements. Nucleic Acids Res 45, 8697–8711 (2017).

33. Lieberman-Aiden, E. et al. Comprehensive mapping of long-range interactions reveals folding principles of the human genome. Science 326, 289–93 (2009).

34. Cairns, J. et al. CHiCAGO: robust detection of DNA looping interactions in Capture Hi-C data. Genome Biol 17, 127 (2016).

35. Cairns, J., Orchard, W.R., Malysheva, V. & Spivakov, M. Chicdiff: a computational pipeline for detecting differential chromosomal interactions in Capture Hi-C data. Bioinformatics 35, 4764–4766 (2019).

36. Subramanian, A. et al. Gene set enrichment analysis: a knowledge-based approach for interpreting genome-wide expression profiles. Proc Natl Acad Sci U S A 102, 15545–50 (2005).

37. Zuin, J. et al. Nonlinear control of transcription through enhancer-promoter interactions. bioRxiv (2021).

38. Zhu, I., Song, W., Ovcharenko, I. & Landsman, D. A model of active transcription hubs that unifies the roles of active promoters and enhancers. Nucleic Acids Res (2021).

39. Fullwood, M.J. et al. An oestrogen-receptor-alpha-bound human chromatin interactome. Nature 462, 58–64 (2009).

40. Nye, A.C. et al. Alteration of large-scale chromatin structure by estrogen receptor. Mol Cell Biol 22, 3437–49 (2002).

41. Rafique, S., Thomas, J.S., Sproul, D. & Bickmore, W.A. Estrogen-induced chromatin decondensation and nuclear re-organization linked to regional epigenetic regulation in breast cancer. Genome Biol 16, 145 (2015).

42. Yin, Y. et al. Impact of cytosine methylation on DNA binding specificities of human transcription factors. Science 356(2017).

43. Gee, J.M. et al. Antihormone induced compensatory signalling in breast cancer: an adverse event in the development of endocrine resistance. Horm Mol Biol Clin Investig 5, 67–77 (2011).

44. Stone, A. et al. DNA methylation of oestrogen-regulated enhancers defines endocrine sensitivity in breast cancer. Nat Commun 6, 7758 (2015).

45. Achinger-Kawecka, J., Chia, K-M., Portman, N., Campbell, E., Du, Q., Laven-Law, G., Clifton, S., Milioli, H.H., Schmitt, A., Wong, E., Hickey, T., Lim, E., Stirzaker, C., Clark, S.J. Epigenetic therapy suppresses tumour growth by rewiring ER-mediated long-range chromatin interactions in ER+ endocrine-resistant breast cancer. bioRxiv (2021).

46. Linnekamp, J.F., Butter, R., Spijker, R., Medema, J.P. & van Laarhoven, H.W.M. Clinical and biological effects of demethylating agents on solid tumours - A systematic review. Cancer Treat Rev 54, 10–23 (2017).

47. Oronsky, B., Oronsky, N., Knox, S., Fanger, G. & Scicinski, J. Episensitization: therapeutic tumor resensitization by epigenetic agents: a review and reassessment. Anticancer Agents Med Chem 14, 1121–7 (2014).

48. Tsai, H.C. et al. Transient Low Doses of DNA-Demethylating Agents Exert Durable Antitumor Effects on Hematological and Epithelial Tumor Cells. Cancer Cell 21, 430–446 (2012).

49. Lu, Z. et al. Epigenetic therapy inhibits metastases by disrupting premetastatic niches. Nature 579, 284–290 (2020).

50. Yu, J. et al. DNA methyltransferase expression in triple-negative breast cancer predicts sensitivity to decitabine. J Clin Invest 128, 2376–2388 (2018).

51. Dahn, M.L. et al. Decitabine Response in Breast Cancer Requires Efficient Drug Processing and Is Not Limited by Multidrug Resistance. Mol Cancer Ther 19, 1110–1122 (2020).

52. de Cubas, A.A. et al. DNA hypomethylation promotes transposable element expression and activation of immune signaling in renal cell cancer. JCI Insight 5(2020).

53. Johnstone, S.E. et al. Large-Scale Topological Changes Restrain Malignant Progression in Colorectal Cancer. Cell 182, 1474–1489 e23 (2020).

54. Buitrago, D. et al. Impact of DNA methylation on 3D genome structure. Nat Commun 12, 3243 (2021).

55. Flavahan, W.A. et al. Altered chromosomal topology drives oncogenic programs in SDH-deficient GISTs. Nature 575, 229–233 (2019).

56. Xu, J. et al. Subtype-specific 3D genome alteration in acute myeloid leukaemia. Nature (2022).

57. Lee, D.S. et al. Simultaneous profiling of 3D genome structure and DNA methylation in single human cells. Nat Methods 16, 999–1006 (2019).

58. Bell, A.C. & Felsenfeld, G. Methylation of a CTCF-dependent boundary controls imprinted expression of the Igf2 gene. Nature 405, 482–5 (2000).

59. Hark, A.T. et al. CTCF mediates methylation-sensitive enhancer-blocking activity at the H19/Igf2 locus. Nature 405, 486–9 (2000).

60. Maurano, M.T. et al. Role of DNA Methylation in Modulating Transcription Factor Occupancy. Cell Rep 12, 1184–95 (2015).

61. Ahmed, M. et al. CRISPRi screens reveal a DNA methylation-mediated 3D genome dependent causal mechanism in prostate cancer. Nat Commun 12, 1781 (2021).

62. Du, Q. et al. DNA methylation is required to maintain DNA replication timing precision and 3D genome integrity. bioRxiv (2020).

63. McLaughlin, K. et al. DNA Methylation Directs Polycomb-Dependent 3D Genome Re-organization in Naive Pluripotency. Cell Rep 29, 1974–1985 e6 (2019).

64. Broome, R. et al. TET2 is a component of the estrogen receptor complex and controls 5mC to 5hmC conversion at estrogen receptor cis-regulatory regions. Cell Rep 34, 108776 (2021).

65. Bi, M. et al. Enhancer reprogramming driven by high-order assemblies of transcription factors promotes phenotypic plasticity and breast cancer endocrine resistance. Nat Cell Biol 22, 701–715 (2020).

66. Nagarajan, S. et al. ARID1A influences HDAC1/BRD4 activity, intrinsic proliferative capacity and breast cancer treatment response. Nat Genet 52, 187–197 (2020).

67. Schmitt, A.D. et al. A Compendium of Chromatin Contact Maps Reveals Spatially Active Regions in the Human Genome. Cell Rep 17, 2042–2059 (2016).

68. Oudelaar, A.M. et al. A revised model for promoter competition based on multiway chromatin interactions at the alpha-globin locus. Nat Commun 10, 5412 (2019).

69. Thomas, H.F. et al. Temporal dissection of an enhancer cluster reveals distinct temporal and functional contributions of individual elements. Mol Cell (2021).

70. DeRose, Y.S. et al. Tumor grafts derived from women with breast cancer authentically reflect tumor pathology, growth, metastasis and disease outcomes. Nature Medicine 17, 1514–U227 (2011).

71. Schoenfelder, S. et al. The pluripotent regulatory circuitry connecting promoters to their long-range interacting elements. Genome Res 25, 582–97 (2015).

72. Schoenfelder, S., Javierre, B.M., Furlan-Magaril, M., Wingett, S.W. & Fraser, P. Promoter Capture Hi-C: High-resolution, Genome-wide Profiling of Promoter Interactions. Jove-Journal of Visualized Experiments (2018).

73. Schoenfelder, S. et al. Polycomb repressive complex PRC1 spatially constrains the mouse embryonic stem cell genome. Nat Genet 47, 1179–1186 (2015).

74. Aryee, M.J. et al. Minfi: a flexible and comprehensive Bioconductor package for the analysis of Infinium DNA methylation microarrays. Bioinformatics 30, 1363–9 (2014).

75. Pidsley, R. et al. Critical evaluation of the Illumina MethylationEPIC BeadChip microarray for whole-genome DNA methylation profiling. Genome Biol 17, 208 (2016).

76. Zhou, W.D., Laird, P.W. & Shen, H. Comprehensive characterization, annotation and innovative use of Infinium DNA methylation BeadChip probes. Nucleic Acids Research 45 (2017).

77. Ritchie, M.E. et al. limma powers differential expression analyses for RNA-sequencing and microarray studies. Nucleic Acids Res 43, e47 (2015).

78. Peters, T.J. et al. De novo identification of differentially methylated regions in the human genome. Epigenetics Chromatin 8, 6 (2015).

79. Wingett, S.W. & Andrews, S. FastQ Screen: A tool for multi-genome mapping and quality control. F1000Res 7, 1338 (2018).

80. Conway, T. et al. Xenome--a tool for classifying reads from xenograft samples. Bioinformatics 28, i172–8 (2012).

81. Dozmorov, M.G. et al. Chromatin conformation capture (Hi-C) sequencing of patient-derived xenografts: analysis guidelines. Gigascience 10(2021).

82. Servant, N. et al. HiC-Pro: an optimized and flexible pipeline for Hi-C data processing. Genome Biology 16(2015).

83. Imakaev, M. et al. Iterative correction of Hi-C data reveals hallmarks of chromosome organization. Nat Methods 9, 999–1003 (2012).

84. Wu, P. et al. 3D genome of multiple myeloma reveals spatial genome disorganization associated with copy number variations. Nat Commun 8, 1937 (2017).

85. Durand, N.C. et al. Juicer Provides a One-Click System for Analyzing Loop-Resolution Hi-C Experiments. Cell Syst 3, 95–8 (2016).

86. Durand, N.C. et al. Juicebox Provides a Visualization System for Hi-C Contact Maps with Unlimited Zoom. Cell Syst 3, 99–101 (2016).

87. Zhou, X. et al. Exploring long-range genome interactions using the WashU Epigenome Browser. Nat Methods 10, 375–6 (2013).

88. Kruse, K., Hug, C.B., Hernandez-Rodriguez, B. & Vaquerizas, J.M. TADtool: visual parameter identification for TAD-calling algorithms. Bioinformatics 32, 3190–3192 (2016).

89. van der Weide, R.H. et al. Hi-C Analyses with GENOVA: a case study with cohesin variants. bioRxiv (2021).

90. Wingett, S. et al. HiCUP: pipeline for mapping and processing Hi-C data. F1000Res 4, 1310 (2015).

91. Blighe K, S.R., M Lewis. EnhancedVolcano: Publication-ready volcano plots with enhanced colouring and labeling. (2018).

92. Dobin, A. et al. STAR: ultrafast universal RNA-seq aligner. Bioinformatics 29, 15–21 (2013).

93. Robinson, M.D. & Oshlack, A. A scaling normalization method for differential expression analysis of RNA-seq data. Genome Biol 11, R25 (2010).

94. Robinson, M.D., McCarthy, D.J. & Smyth, G.K. edgeR: a Bioconductor package for differential expression analysis of digital gene expression data. Bioinformatics 26, 139–40 (2010).

95. Jin, Y., Tam, O.H., Paniagua, E. & Hammell, M. TEtranscripts: a package for including transposable elements in differential expression analysis of RNA-seq datasets. Bioinformatics 31, 3593–3599 (2015).

96. Love, M.I., Huber, W. & Anders, S. Moderated estimation of fold change and dispersion for RNA-seq data with DESeq2. Genome Biology 15(2014).

97. Langmead, B., Trapnell, C., Pop, M. & Salzberg, S.L. Ultrafast and memory-efficient alignment of short DNA sequences to the human genome. Genome Biol 10, R25 (2009).

98. Zhang, Y. et al. Model-based analysis of ChIP-Seq (MACS). Genome Biol 9, R137 (2008).

99. Feng, J.X., Liu, T., Qin, B., Zhang, Y. & Liu, X.S. Identifying ChIP-seq enrichment using MACS. Nature Protocols 7, 1728–1740 (2012).

100. Heger, A., Webber, C., Goodson, M., Ponting, C.P. & Lunter, G. GAT: a simulation framework for testing the association of genomic intervals. Bioinformatics 29, 2046–8 (2013).

101. Yu, G., Wang, L.G. & He, Q.Y. ChIPseeker: an R/Bioconductor package for ChIP peak annotation, comparison and visualization. Bioinformatics 31, 2382–3 (2015).

102. Ramirez, F. et al. deepTools2: a next generation web server for deep-sequencing data analysis. Nucleic Acids Res 44, W160–5 (2016).

103. Heinz, S. et al. Simple Combinations of Lineage-Determining Transcription Factors Prime cis-Regulatory Elements Required for Macrophage and B Cell Identities. Molecular Cell 38, 576–589 (2010).

