## Supplementary Notre for "Epigenetic therapy targets the 3D epigenome in endocrine-resistant breast cancer"

**Supplementary Notes**

**Decitabine-induced DNA hypomethylation activates transposable element (TE) expression**

Epigenetic therapies that target repressive epigenetic machinery have also been reported to mediate their anti-tumour effect by activating transposable elements (TEs) *via* the viral mimicry response[1,2]. Therefore to study the viral mimicry response to Decitabine in our models, we first evaluated genome-wide methylation levels at different classes of repetitive elements using the REMP package[3], including LTRs, LINE1 elements and Alu elements. We observed genome-wide loss of DNA methylation at all repeat element sub-groups (Supplementary Fig. 1a, b). All repeat element classes showed ~12% loss of DNA methylation in Decitabine treated tumours. Notably, although all TE classes showed some degree of DNA methylation loss, the extent of DNA hypomethylation measured at repeat elements was less than those observed genome-wide and significantly less to DNA hypomethylation at enhancer regions (Extended Data Fig. 1h).

To assess whether Decitabine-induced hypomethylation increased transcription of TEs, we quantified global TE expression in poly-A enriched RNA-seq data using TEtranscripts[4] in four replicates of Vehicle and Decitabine-treated tumours. We detected a total of ~1079 TE transcripts in our samples (1049 in Gar15-13 and 1110 in HCI-005). Differential analysis of TE expression (DESeq2) revealed an increase in TE expression in both PDX models following treatment with Decitabine (Supplementary Fig. 1c, d), in agreement with previous studies showing that DNA hypomethylation activates TE expression in other model systems[5]. In total, 116 and 68 TEs were induced in Decitabine-treated Gar15-13 and HCI-005 tumours, respectively (logFC > 0.2 and FDR < 0.1) (Supplementary Fig. 1c, d). In both PDX models, LTRs and LINE1 elements were the most abundant class of differentially expressed TEs, followed by DNA transposons (Supplementary Fig. 1e, f).

To assess how loss of DNA methylation affected related antiviral signalling pathways in the PDX tumours, we investigated gene expression changes beyond TEs in Gar15-

13 PDX tumours. Gene set enrichment analysis (GSEA) revealed that many immune pathways (inflammatory response, interferon response, IL6 signalling) were among the most upregulated in comparing Decitabine and Vehicle treated samples. Genes belonging to these pathways were up-regulated in Decitabine-treated tumours, including canonical components of the antiviral pathway (for example *CDKN1A*, *HIF1A* or *IFI27*) (Supplementary Fig. 1g, h), consistent with antiviral signalling induced by Decitabine.

**Multi-omics analyses of ER+ breast cancer PDX tumours following Decitabine treatment**

To provide a comprehensive characterisation of molecular changes following epigenetic therapy we used a multi-omics approach to generate and interrogate genome-wide maps of PDX breast tumours treated with and without Decitabine. We performed *in situ* Hi-C, promoter-anchored interactome (Promoter Capture Hi-C (PCHi-C)), DNA methylation (EPIC Microarrays), ER and FOXA1 cistromes (ChIP-seq) and gene expression profiles (RNA-seq) in the same tumours. Together, we generated 47 libraries and more than 5 billion sequencing reads for Hi-C (Supplementary Table 2), PCHi-C (Supplementary Table 3), RNA-seq (Supplementary Table 4) and ChIP-seq (Supplementary Table 5).

**Unsupervised clustering of multi-omics data**

We initially explored the inter-subpopulation variability across the multi-omics datasets in the two PDX models (Gar15-13 and HCI-005) to determine if Decitabine treatment resulted in changes to high-level genome architecture. By comparing the top 10,000 most variable interactions between the Hi-C maps, we observed clear difference between Vehicle and Decitabine-treated tumours (Supplementary Fig. 2a), suggesting large-scale changes in the overall genome architecture. Furthermore, we interrogated Hi-C data from two endocrine-sensitive MCF7 cell lines and MCF7 tamoxifen-resistant derivative (TAMR) cell lines from [6] as well as the MCF7 cell line from [7] and performed MDS analysis. The comparison of samples shows that Decitabine treatment induced a 3D genome organisation that is more similar to endocrine-sensitive MCF7 cells and clearly different from Vehicle-treated tumours, which clustered more closely to the tamoxifen-resistant TAMR cell line (Supplementary Fig. 2a). Principal component analyses (PCA) of DNA methylation (Supplementary Fig. 2b), ER cistromes (Supplementary Fig. 2c) and gene expression profiles (Supplementary Fig. 2d) further confirmed large-scale differences were also induced in the methylome, ER binding and expression by Decitabine treatment.

**Quality control of Hi-C data**

The generated Hi-C data was of high quality in the low-input, snap-frozen PDX tumour tissues, as characterised by the high percentage of valid interactions (~90%), high cis/trans ratio (~80:20%) and high percentage of long-range interactions (60% >10KBs) (Supplementary Table 2 and Supplementary Fig. 3a). Host mouse reads were quantified (Supplementary Fig. 3b) and removed as described in [8]. We observed strong concordance between Hi-C replicates as measured by the HiCRep SCC score [9] (Supplementary Fig. 3c).

**Relationship between differential TADs and A/B compartment switching**

The majority of differential TAD boundaries were located within stable A-type or B-type compartments (>95%) (Supplementary Fig. 3d), while a small number of lost TAD boundaries were located at regions that shifted from B-type to A-type assignment (81 TAD boundaries; Supplementary Fig. 3d).

**Detection of differential promoter-anchored interactions from PCHi-C data**

Promoter Capture Hi-C (PCHi-C) was performed as described previously [10-12]. Hi-C libraries were hybridised to custom-designed genomic restriction fragments covering 23,711 annotated gene promoters. We generated ~740,000,000 total valid read-pairs (di-tags) from the three PDX tumours for each treatment group, obtaining ~125 million valid ligation products per sample (~350 – 400 million per treatment) (Supplementary Table 3). To identify statistically significant interactions between promoters and other regulatory elements from the PCHi-C data, we used the CHiCAGO pipeline [13]. We obtained on average >100,000 statistically significant interactions per replicate. Identified interactions were highly enriched for histone modifications associated with open chromatin and regulatory elements (publicly available ChIP-seq datasets: H3K4me3, H3K4me1 and H3K27ac) as compared to random regions (Supplementary Fig. 4a, b).

To identify differential promoter-anchored interactions, we first used Chicdiff to calculate the asinh-transformed CHiCAGO scores and log fold changes between Vehicle and Decitabine tumours for each CHiCAGO interaction. We next integrated the Chicdiff scores with an intersection of CHiCAGO interactions identified in Vehicle and in Decitabine tumours and defined gained, maintained and lost interacting regions separated into promoter baits and enhancer OEs.

**Functional studies of DNA hypomethylation and re-methylation in TAMR cells**

First, to determine appropriate Decitabine dose, cytotoxicity was assessed using alamarBlue assay and 100nM dose was selected for subsequent experiments. Tamoxifen-resistant (TAMR) MCF7 cells were seeded in a white flat-bottom 96-well plate at a density of 2500 cells/well based on doubling time and allowed to adhere overnight. Cells were treated daily with Decitabine (0nM, 10nM, 50nM, 100nM, 200nM, 300nM, 500nM, 1uM, 5uM, 10uM and 100uM) for 72h and cellular viability was assessed on day 5, following 2 days of recovery using the alamarBlue (ThermoFisher, DAL1025) assay in accordance with the manufacture's recommendations. The normalised fluorescence intensity (A570/EM590) was calculated by subtracting the average background fluorescence intensity detected in wells containing media + 10% alamarBlue and no cells from the fluorescence intensity detected in wells containing cells. Based on cellular viability assayed, 100nM Decitabine dose was selected for further experiments (Supplementary Fig. 5a). Cells were then treated daily with decitabine (100 nM) for 7 consecutive days (Day 7 Decitabine samples). For Decitabine Recovery samples, after 7 days of Decitabine treatment, fresh media was added, and cells were cultured for 28 additional days and harvested on day 35. Control cells were collected at 11 days (Control Early) and 35 days (Control Late). Protein, DNA and RNA were harvested for Day 7 Decitabine, Decitabine Recovery and Control Early and Late samples. Loss and recovery of DNMT1 protein expression was confirmed by Western blot (Supplementary Fig. 5b).

***Protein extraction and immunoblot analysis.*** Whole cell protein lysates were prepared by resuspending scaped cells in modified RIPA buffer (50 mM Tris-HCL pH 7.5, 150 mM NaCl, 1 mM EDTA, 1 mM NaF, 1% Igepal, 0.25% Sodium-deoxycholate) with protease inhibitors and incubated on ice for 60 min with vortexing every 10 min. Lysate was sonicated using the Bioruptor on the high setting, 6x cycles of 30sec on/

30sec off and stored at −80°C. Protein concentration was determined using a BCA assay (Pierce, #23227) according to manufacturer's instructions. Protein lysate was prepared using NuPAGE® LDS sample buffer (Life Technologies, NP0007) and NuPAGE® sample reducing agent (Life Technologies, NP0004) followed by heating at 95°C for 5 min. Samples were resolved by gel electrophoresis using the NuPage Bis-Tris 4%–12% precast gel system according to the manufacturer's instructions (Life Technologies). Western blot transfer was carried out according to the manufacturer's instructions with the SureLock X-cell system (ThermoFisher Scientific) using transfer buffer containing 20% methanol content. Antibody used was C-terminal DNMT1 – Abcam #ab92314; GAPDH – Invitrogen #AM4300. Western blots (WBs) were then treated with Western Lightning Plus-ECL (Perkin Elmer, #NEL104001EA) before developing on Super Rx Fuji Medical X-Ray Film (Fujifilm, #4741019289) using the Konica Tabletop X-Ray Film Processor (#SRX101A). Developed film was scanned using the Epson Perfection V800/850 scanner.

**Supplementary References**


1. Roulois, D. *et al.* DNA-Demethylating Agents Target Colorectal Cancer Cells by Inducing Viral Mimicry by Endogenous Transcripts. *Cell* **162**, 961-973 (2015).
2. Mehdipour, P. *et al.* Epigenetic therapy induces transcription of inverted SINEs and ADAR1 dependency. *Nature* **588**, 169-+ (2020).
3. Zheng, Y. *et al.* Prediction of genome-wide DNA methylation in repetitive elements. *Nucleic Acids Res* **45**, 8697-8711 (2017).
4. Jin, Y., Tam, O.H., Paniagua, E. & Hammell, M. TEtranscripts: a package for including transposable elements in differential expression analysis of RNA-seq datasets. *Bioinformatics* **31**, 3593-3599 (2015).
5. de Cubas, A.A. *et al.* DNA hypomethylation promotes transposable element expression and activation of immune signaling in renal cell cancer. *JCI Insight* **5**(2020).
6. Achinger-Kawecka, J. *et al.* Epigenetic reprogramming at estrogen-receptor binding sites alters 3D chromatin landscape in endocrine-resistant breast cancer. *Nat Commun* **11**, 320 (2020).
7. Barutcu, A.R. *et al.* Chromatin interaction analysis reveals changes in small chromosome and telomere clustering between epithelial and breast cancer cells. *Genome Biol* **16**, 214 (2015).
8. Dozmorov, M.G. *et al.* Chromatin conformation capture (Hi-C) sequencing of patient-derived xenografts: analysis guidelines. *Gigascience* **10**(2021).
9. Yang, T. *et al.* HiCRep: assessing the reproducibility of Hi-C data using a stratum-adjusted correlation coefficient. *Genome Research* **27**, 1939-1949 (2017).
10. Schoenfelder, S. & Fraser, P. Long-range enhancer-promoter contacts in gene expression control. *Nat Rev Genet* **20**, 437-455 (2019).

11. Schoenfelder, S., Javierre, B.M., Furlan-Magaril, M., Wingett, S.W. & Fraser, P. Promoter Capture Hi-C: High-resolution, Genome-wide Profiling of Promoter Interactions. *Jove-Journal of Visualized Experiments* (2018).
12. Jung, I. *et al.* A compendium of promoter-centered long-range chromatin interactions in the human genome. *Nat Genet* **51**, 1442-1449 (2019).
13. Cairns, J. *et al.* CHiCAGO: robust detection of DNA looping interactions in Capture Hi-C data. *Genome Biol* **17**, 127 (2016).

### Supplementary Figure 1

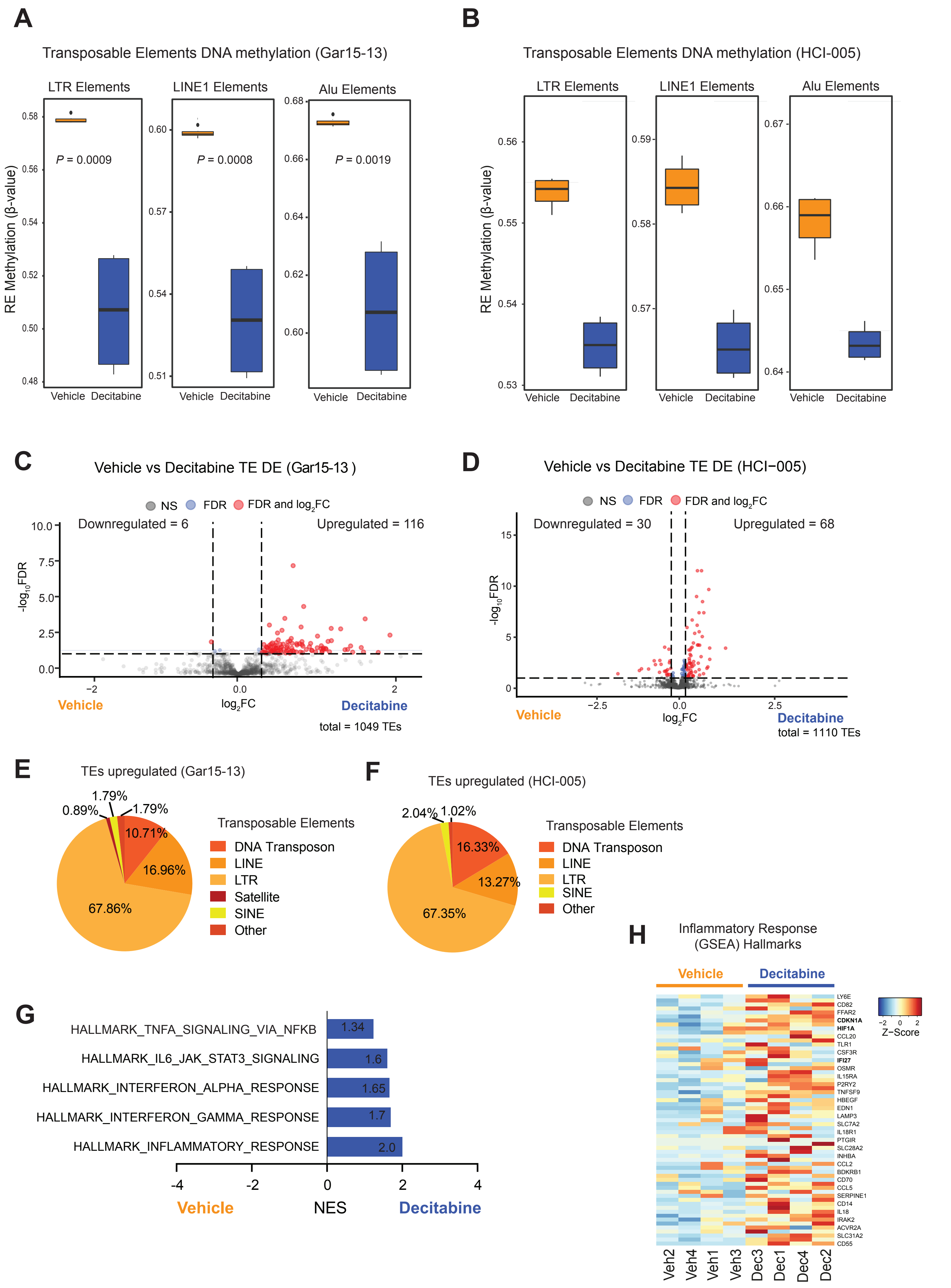

**Supplementary Fig. 1**

**(a)** Boxplots showing the distribution of DNA methylation for Gar15-13 Vehicle and Decitabine-treated tumours for EPIC probes mapping to LTR, LINE1 and Alu repetitive elements (REMP annotation). Black line indicates mean ± SD.

**(b)** Boxplots showing the distribution of DNA methylation profiles for HCI-005 Vehicle and Decitabine-treated tumours for EPIC probes mapping to LTR, LINE1 and Alu repetitive elements (REMP annotation). Black line indicates mean ± SD.

**(c)** Volcano plot showing the differential expression of transposable elements in Decitabine-treated Gar15-13 tumours compared to Vehicle. Lines indicate differentially expressed TEs considered significant (logFC > 0.2 and FDR < 0.1).

**(d)** Volcano plot showing the differential expression of transposable elements in Decitabine-treated HCI-005 tumours as compared to Vehicle. Lines indicate differentially expressed TEs considered significant (logFC > 0.2 and FDR < 0.1).

**(e)** Pie chart showing distribution of differentially expressed Transposable Elements (TE) classes in Gar15-13.

**(f)** Pie chart showing distribution of differentially expressed TE classes in HCI-005.

**(g)** Normalised enrichment scores (NES) for significantly enriched gene sets belonging to the immune GSEA hallmark in Gar15-13.

**(h)** RNA-seq heatmap of Decitabine-induced changes in expression of genes belonging to the Inflammatory Response (GSEA) Hallmarks in Gar15-13. Top differentially expressed genes plotted (FDR < 0.05).

### Supplementary Figure 2

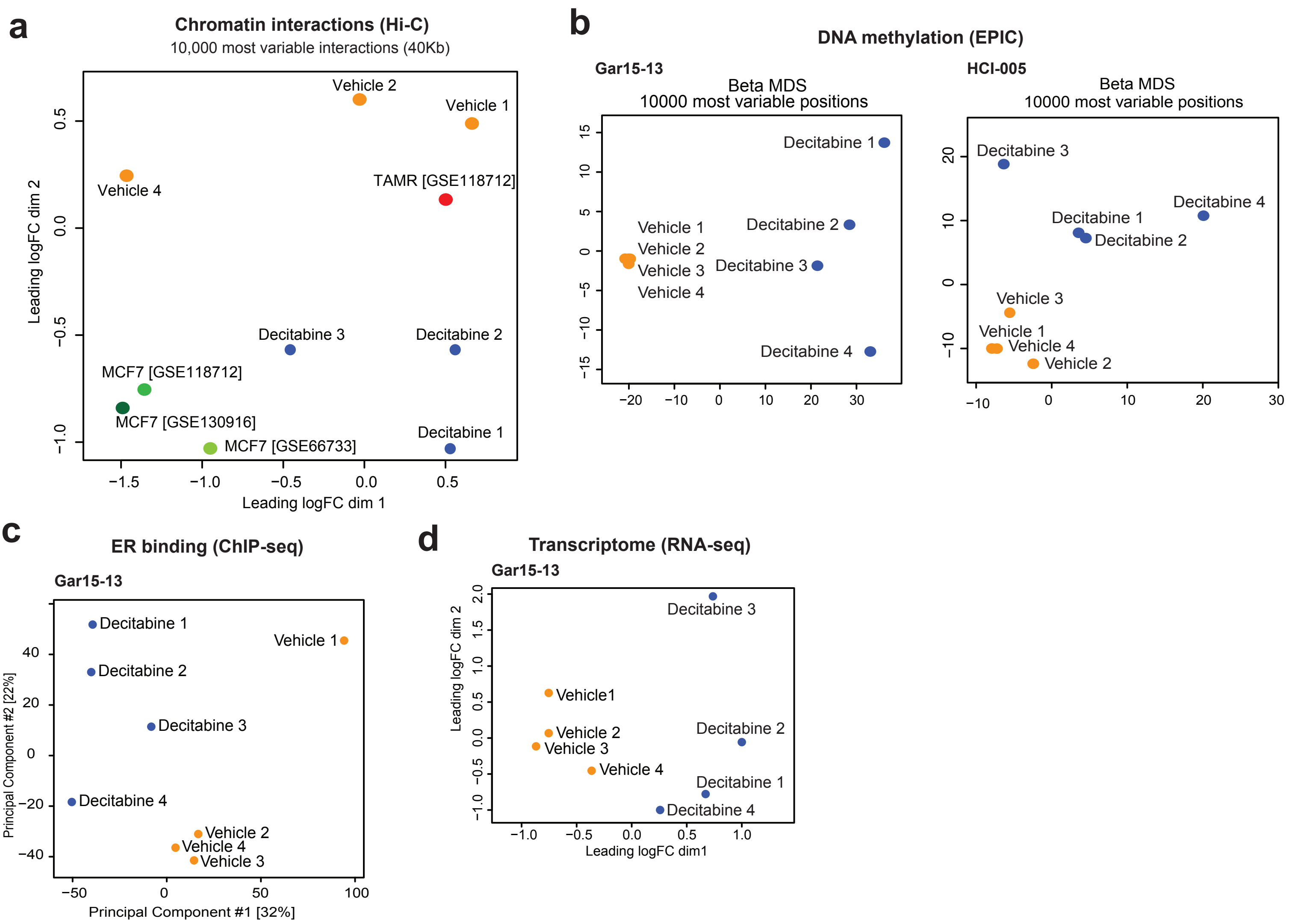

**Supplementary Fig. 2**

**(a)** Multidimensional-scaling plot showing the relationships among the chromatin interaction profiles of Gar15-13 Vehicle tumours (n = 3), Decitabine-treated tumours (n = 3), tamoxifen-resistant MCF7 cells (TAMR; n = 1) and parental MCF7 cells (MCF7; n = 2) and MCF7 cell lines from [7]. Distances on the plot represent the leading log fold change (logFC). Data plotted for top 1000 most variable interactions.

**(b)** Principal component analysis showing the relationship among the DNA methylation profiles of Gar15-13 (left) and HCI-005 (right) Vehicle tumours (n = 4), Decitabine-treated tumours (n = 4). Data plotted for Beta values from top 10,000 most variable probes.

**(c)** Principal component analysis showing the relationship among the ER binding profiles of Gar15-13 (left) Vehicle tumours (n = 4), Decitabine-treated tumours (n = 4). Data plotted for normalised read counts.

**(d)** Principal component analysis showing the relationship among the transcriptome profiles of Gar15-13 (left) Vehicle tumours (n = 4), Decitabine-treated tumours (n = 4). Data plotted for normalised TPM values.

### Supplementary Figure 3

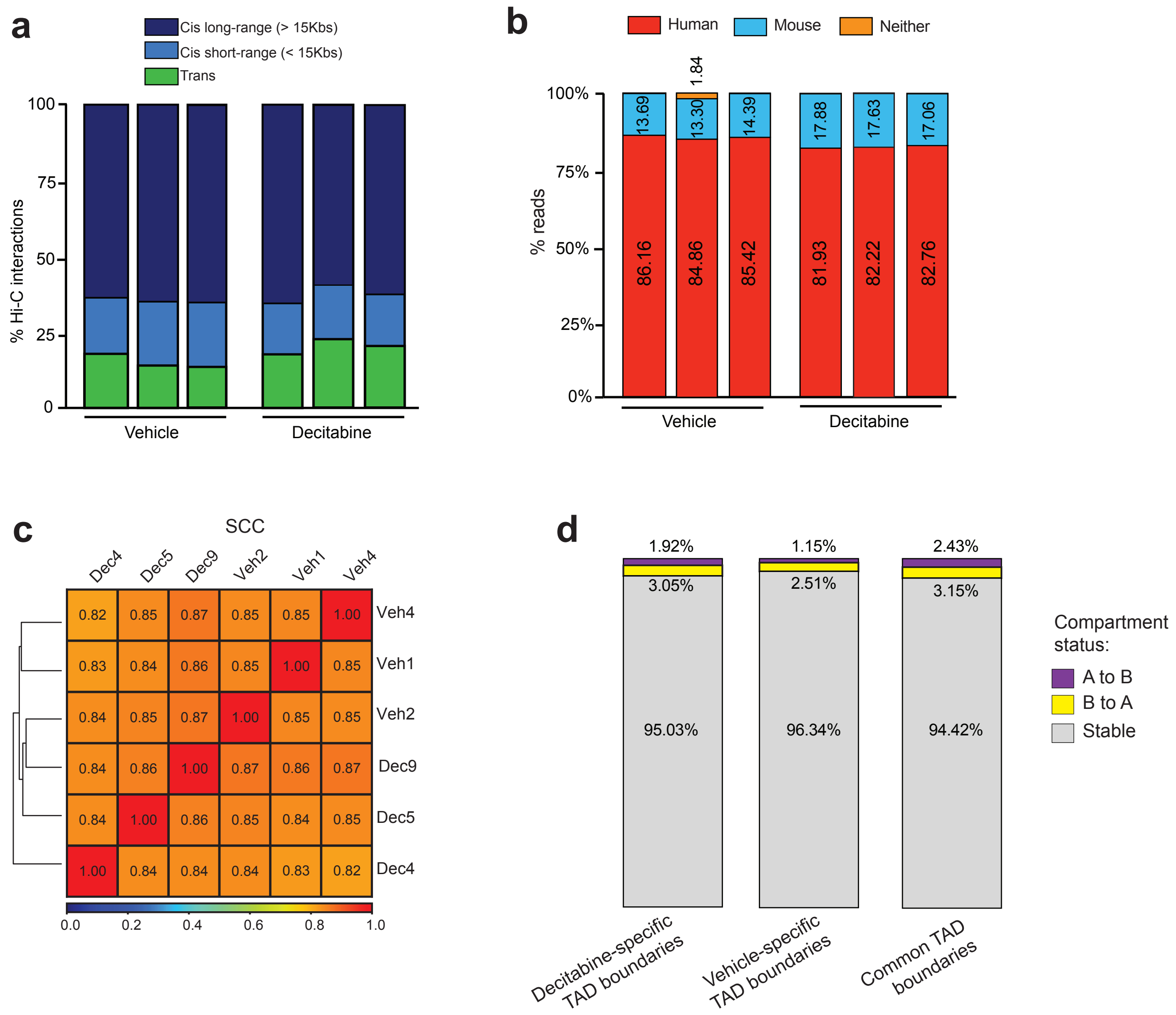

**Supplementary Fig. 3**

**(a)** Barplot showing distribution of short- and long-range intra-chromosomal and inter-chromosomal interactions identified in *in situ* Hi-C data from Gar15-13 PDX tumours.

**(b)** Proportion of human (hg38) and mouse (mm10) reads detected by Xenome in Hi-C data from PDX tumours.

**(c)** Heatmap of stratum adjusted correlation coefficient (SCC) values calculated by HiCRep between the Hi-C interaction maps.

**(d)** Boxplot showing percentage overlap between differential (Decitabine-specific and Vehicle-specific) and common TAD boundaries and A/B compartment switching.

### Supplementary Figure 4

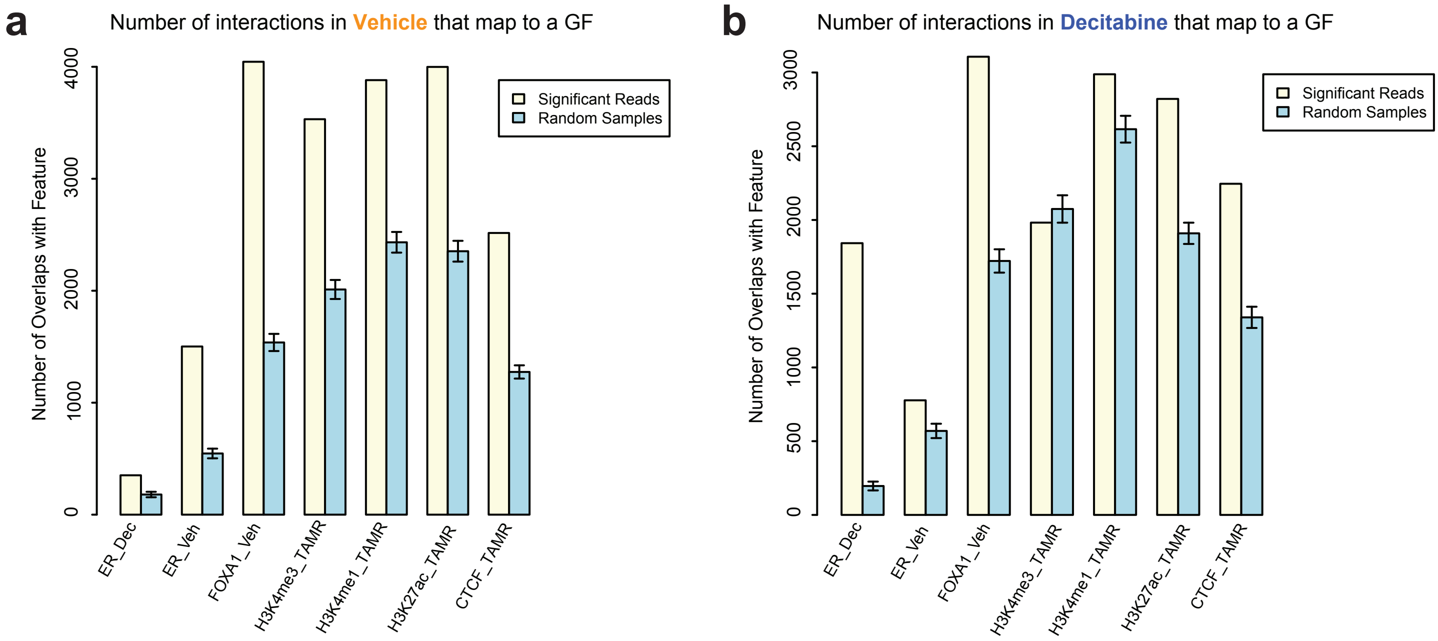

**Supplementary Fig. 4**

**(a)** Significant CHiCAGO promoter-anchored interactions in three Vehicle tumours enrichment for histone marks and transcription factors compared with distance-matched random regions. Error bars show SD across 100 draws of random regions.

**(b)** Significant CHiCAGO promoter-anchored interactions in three Decitabine-treated tumours enrichment for histone marks and transcription factors compared with distance-matched random regions. Error bars show SD across 100 draws of random regions.

### Supplementary Figure 5

a

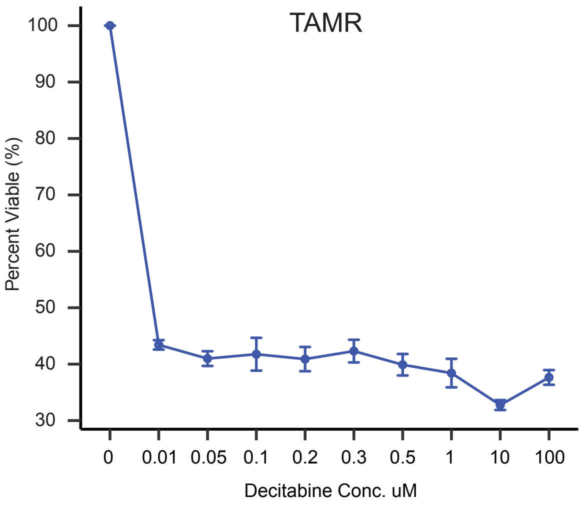


b

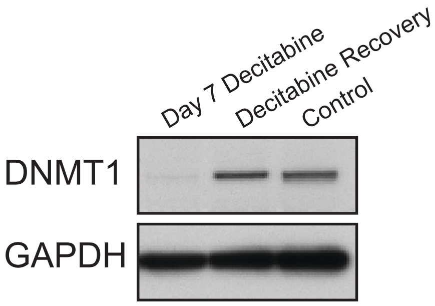

**Supplementary Fig. 5**

**(a)** Endocrine-resistant MCF7 cells (TAMRs) were treated with the indicated doses of decitabine for 3 consecutive days and assayed on day 5. Decitabine dose-response curve for viability in TAMR cells. All data are mean ± SD ($n$ = 4). **(b)** DNMT1 protein levels were assessed in TAMR cells at day 7 of Decitabine treatment (100nM) by immunoblot analysis. GAPDH was included as a loading control.
