## Extended Data Figures for "Epigenetic therapy targets the 3D epigenome in endocrine-resistant breast cancer"

### Extended Data Fig. 1

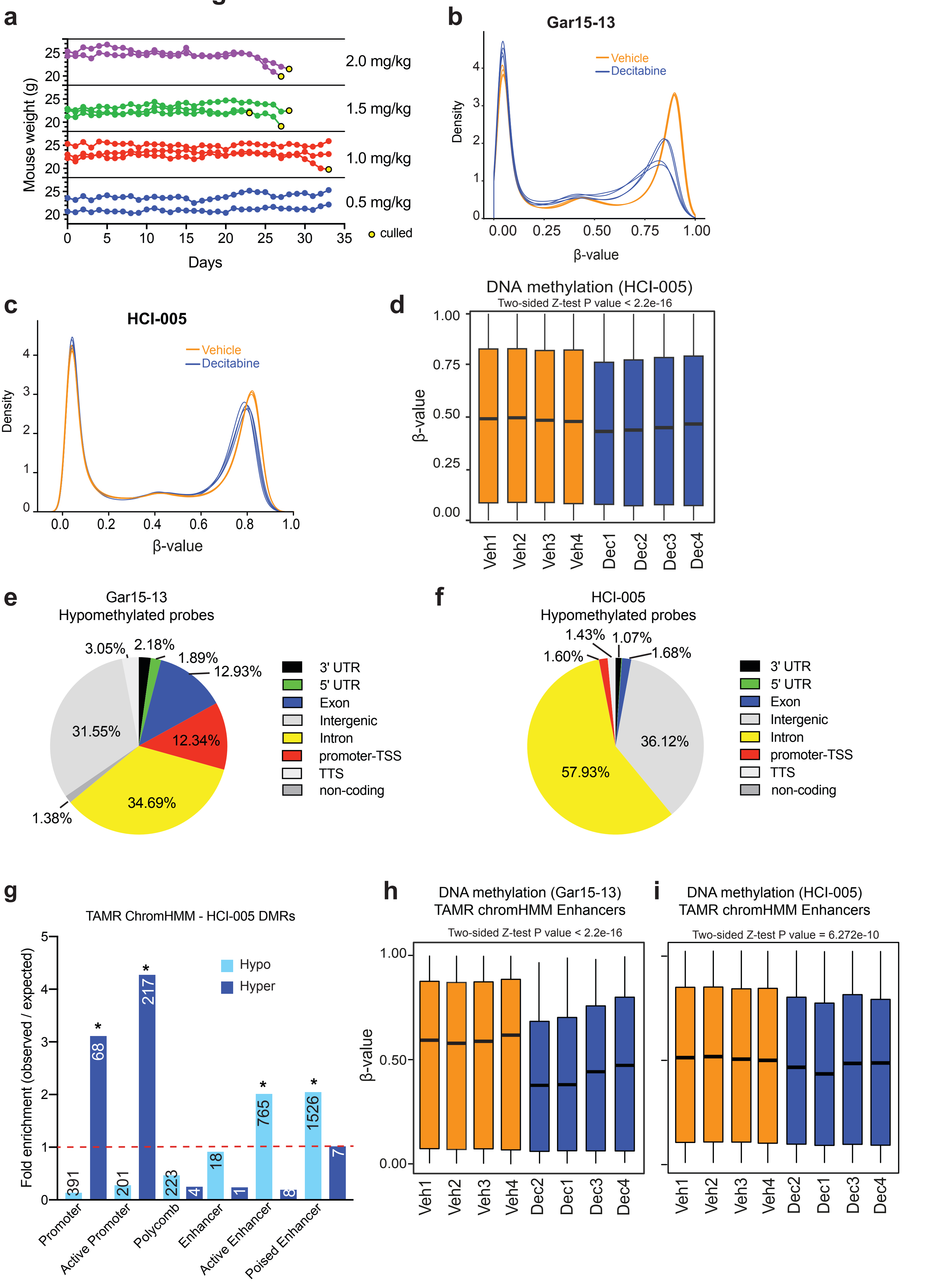

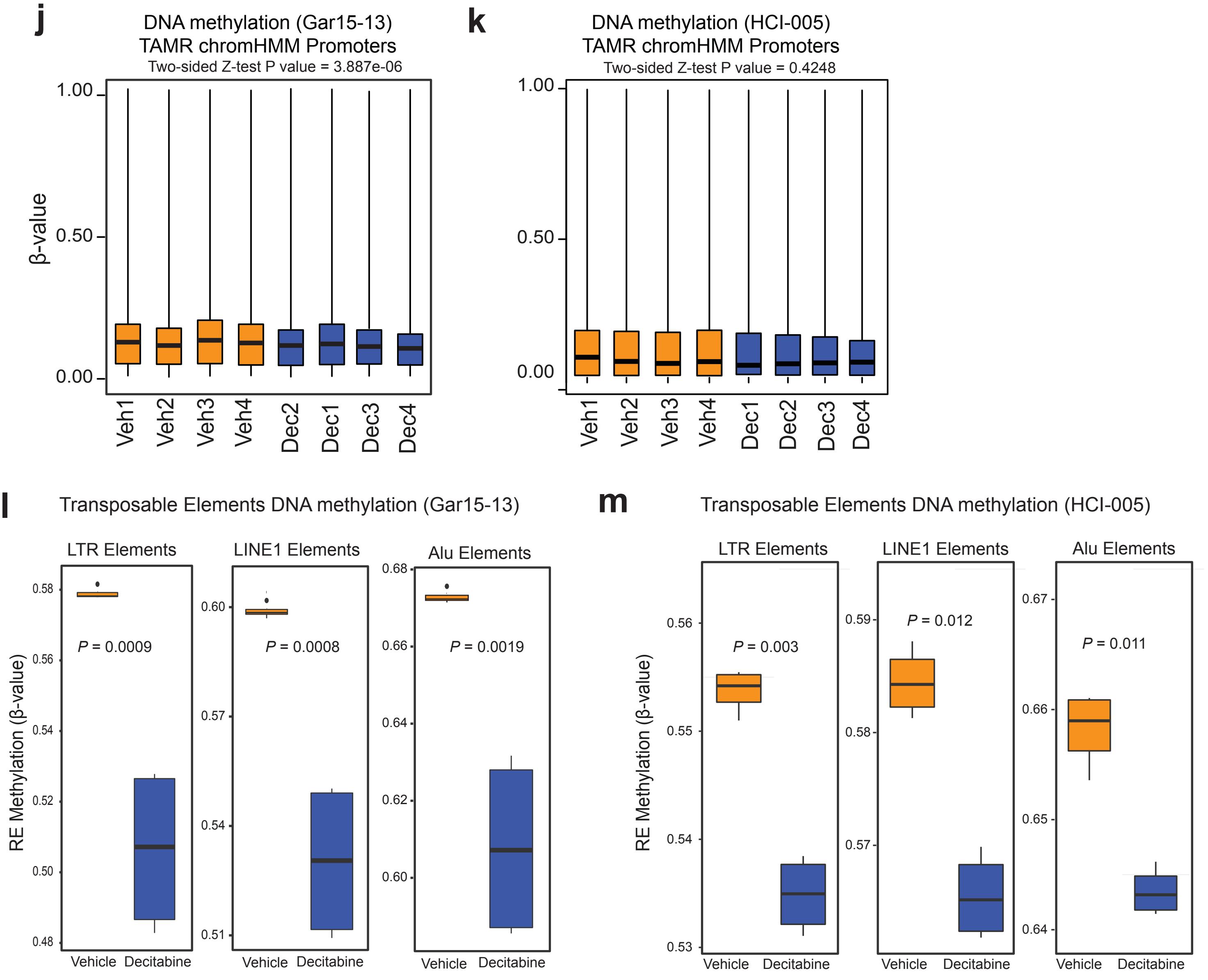

j
DNA methylation (Gar15-13)
TAMR chromHMM Promoters
Two-sided Z-test P value = 3.887e-06
β-value
1.00
0.50
0.00
Veh1
Veh2
Veh3
Veh4
Dec2
Dec1
Dec3
Dec4
k
DNA methylation (HCI-005)
TAMR chromHMM Promoters
Two-sided Z-test P value = 0.4248
1.00
0.50
0.00
Veh1
Veh2
Veh3
Veh4
Dec1
Dec2
Dec3
Dec4
l
Transposable Elements DNA methylation (Gar15-13)
LTR Elements
LINE1 Elements
Alu Elements
RE Methylation (β-value)
P = 0.0009
P = 0.0008
P = 0.0019
Vehicle
Decitabine
m
Transposable Elements DNA methylation (HCI-005)
LTR Elements
LINE1 Elements
Alu Elements
RE Methylation (β-value)
P = 0.003
P = 0.012
P = 0.011
Vehicle
Decitabine

**Extended Data Fig. 1**

**(a)** Mice were treated with indicated doses of Decitabine for 35 consecutive days and mice weight was assayed to determine the most appropriate Decitabine concentration.

**(b)** Density plot showing DNA methylation distribution in four Vehicle and four Decitabine-treated Gar15-13 tumours.

**(c)** Density plot showing DNA methylation distribution in four Vehicle and four Decitabine-treated HCI-005 tumours.

**(d)** Boxplots showing the distribution of DNA methylation profiles for HCI-005 Vehicle and Decitabine-treated tumours for all DNA methylation EPIC probes across the genome.

**(e)** RefSeq annotation of Vehicle vs. Decitabine hypomethylated probes (Gar15-13)

**(f)** RefSeq annotation of Vehicle vs. Decitabine hypomethylated probes (HCI-005)

**(g)** Bar plot showing the association of differentially methylated probes in HCI-005 Decitabine treatment compared to Vehicle across regulatory regions of the genome as determined by TAMR ChromHMM annotation. Observed over expected fold change enrichment shown. * P value < 0.001. The numbers of hypo/hypermethylated probes located within each specific regions are presented in the respective column.

**(h)** Boxplots showing the distribution of DNA methylation profiles for Gar15-13 Vehicle and Decitabine-treated tumours for EPIC probes located at TAMR ChromHMM enhancer regions. Black line indicates median ± SD.

**(i)** Boxplots showing the distribution of DNA methylation profiles for HCI-005 Vehicle and Decitabine-treated tumours for EPIC probes located at TAMR ChromHMM enhancer regions. Black line indicates median ± SD.

**(j)** Boxplots showing the distribution of DNA methylation profiles for Gar15-13 Vehicle and Decitabine-treated tumours for EPIC probes located at TAMR ChromHMM promoter regions. Black line indicates median ± SD.

**(k)** Boxplots showing the distribution of DNA methylation profiles for HCI-005 Vehicle and Decitabine-treated tumours for EPIC probes located at TAMR ChromHMM promoter regions. Black line indicates median ± SD.

**(l)** Observed over expected fold change enrichment of Gar15-13 and HCI-005 Decitabine *vs.* Vehicle hypomethylated DMRs at ER-enhancers hypermethylated with endocrine resistance (Stone *et al.*, 2015). * *P* value < 0.001, n = 1000 permutations.

**(m)** Boxplots showing the distribution of DNA methylation for Gar15-13 Vehicle and Decitabine-treated tumours for EPIC probes mapping to LTR, LINE1 and Alu repetitive elements (REMP annotation). Black line indicates median ± SD.

**(n)** Boxplots showing the distribution of DNA methylation profiles for HCI-005 Vehicle and Decitabine -treated tumours for EPIC probes mapping to LTR, LINE1 and Alu repetitive elements (REMP annotation). Black line indicates median ± SD.

#### Extended Data Fig. 2

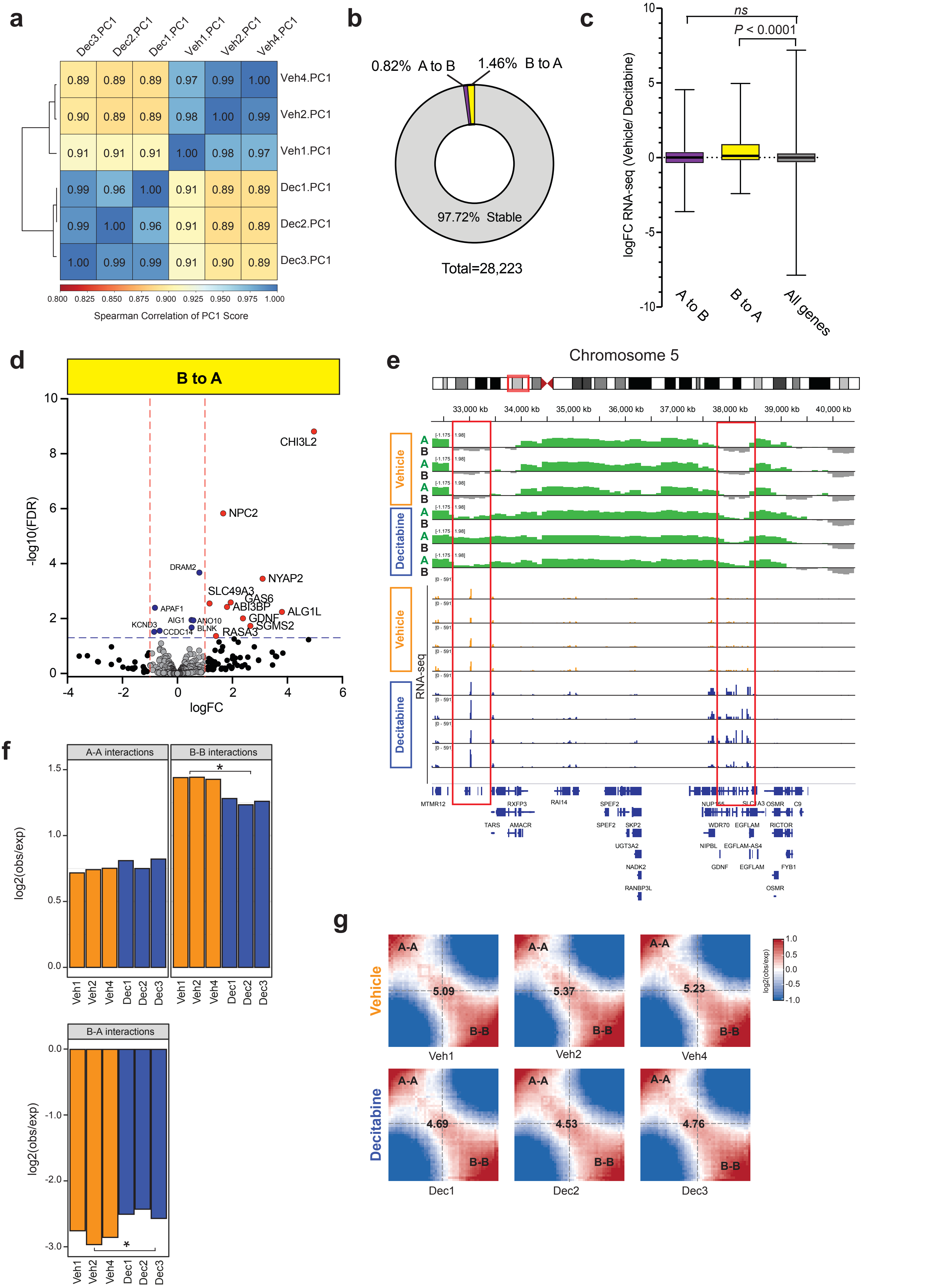

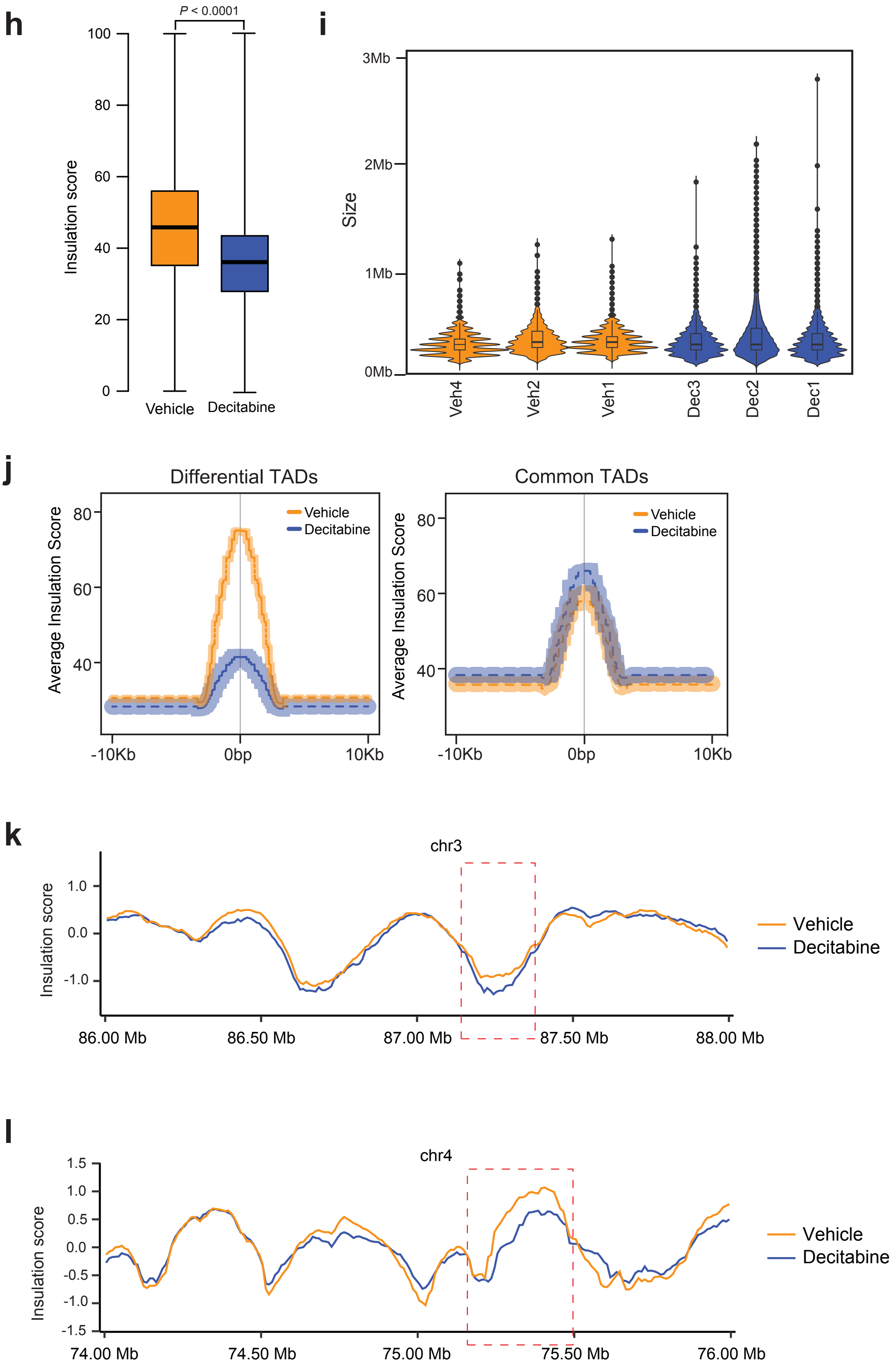
h
P < 0.0001
Insulation score
100
80
60
40
20
0
Vehicle
Decitabine
i
Size
3Mb
2Mb
1Mb
0Mb
Veh4
Veh2
Veh1
Dec3
Dec2
Dec1
j
Differential TADs
Common TADs
Average Insulation Score
80
60
40
Vehicle
Decitabine
-10Kb
0bp
10Kb
k
chr3
Insulation score
1.0
0.0
-1.0
86.00 Mb
86.50 Mb
87.00 Mb
87.50 Mb
88.00 Mb
Vehicle
Decitabine
l
chr4
Insulation score
1.5
1.0
0.5
0.0
-0.5
-1.0
-1.5
74.00 Mb
74.50 Mb
75.00 Mb
75.50 Mb
76.00 Mb
Vehicle
Decitabine

**Extended Data Fig. 2**

**(a)** Heatmap showing Spearman pairwise correlations between the eigenvectors (PC1) in three Decitabine-treated and three Vehicle PDX Gar15-13 tumours. Samples are ordered according to complete linkage hierarchal clustering.

**(b)** Pie chart showing distributing of stable (A to A; B to B) and switching (A to B; B to A) compartments in Decitabine-treated tumours compared to Vehicle

**(c)** Barplot showing logFC expression between Vehicle and Decitabine-treated tumours of genes located either at A to B or B to A switching compartments. *P* value: Wilcox rank-sum test.

**(d)** Volcano plot showing Decitabine *vs.* Vehicle differential expression of genes located at compartment that switched from B to A assignment in Decitabine-treated tumours.

**(e)** Browser snapshot of Hi-C eigenvectors and RNA-seq in Vehicle and Decitabine-treated tumours (n = 3 Hi-C and n = 4 RNA-seq each), showing demarcation of open (A-type; positive values) and closed (B-type; negative values) compartment changes across a region on chromosome 5, which is associated with increased expression of genes located within this region.

**(f)** Barplot showing log2 observed over expected A – A, B – B and B – A compartment interactions in Vehicle and Decitabine tumours. * *P* value two-tailed Students t-test < 0.05

(**g**) Average contact enrichment between pairs of 50Kb loci arranged by their PC1 eigenvector in Vehicle and Decitabine-treated tumours. The numbers at the center of the heatmaps indicate compartment strength calculated as the log2 ratio of (A–A + B–B)/ (A–B + B–A) using the mean values.

**(h)** Boxplot of average insulation score calculated by TADtool at 50Kb resolution in three replicates of Decitabine-treated and Vehicle PDX tumours. *P* value Wilcox rank-sum test.

**(i)** Violin plot showing the distribution in TAD sizes for Vehicle and Decitabine treated Gar15-13 tumours (n = 3 each).

**(j)** Average insulation score at differential and common TADs. Lines show mean values, while light shading represents SEM.

**(k)** Snapshot of region on chromosome 3, showing insulation score calculated in Vehicle and Decitabine-treated tumour Hi-C matrixes, demonstrating loss of TAD boundary insulation is Decitabine-treated samples (indicated with a red box). Merged Hi-C data from replicates shown at 10Kb resolution.

**(l)** Snapshot of region on chromosome 4, showing insulation score calculated in Vehicle and Decitabine-treated tumour Hi-C matrixes, demonstrating loss of TAD boundary insulation is Decitabine-treated samples (indicated with a red box). Merged Hi-C data from replicates shown at 10Kb resolution.

#### Extended Data Fig. 3

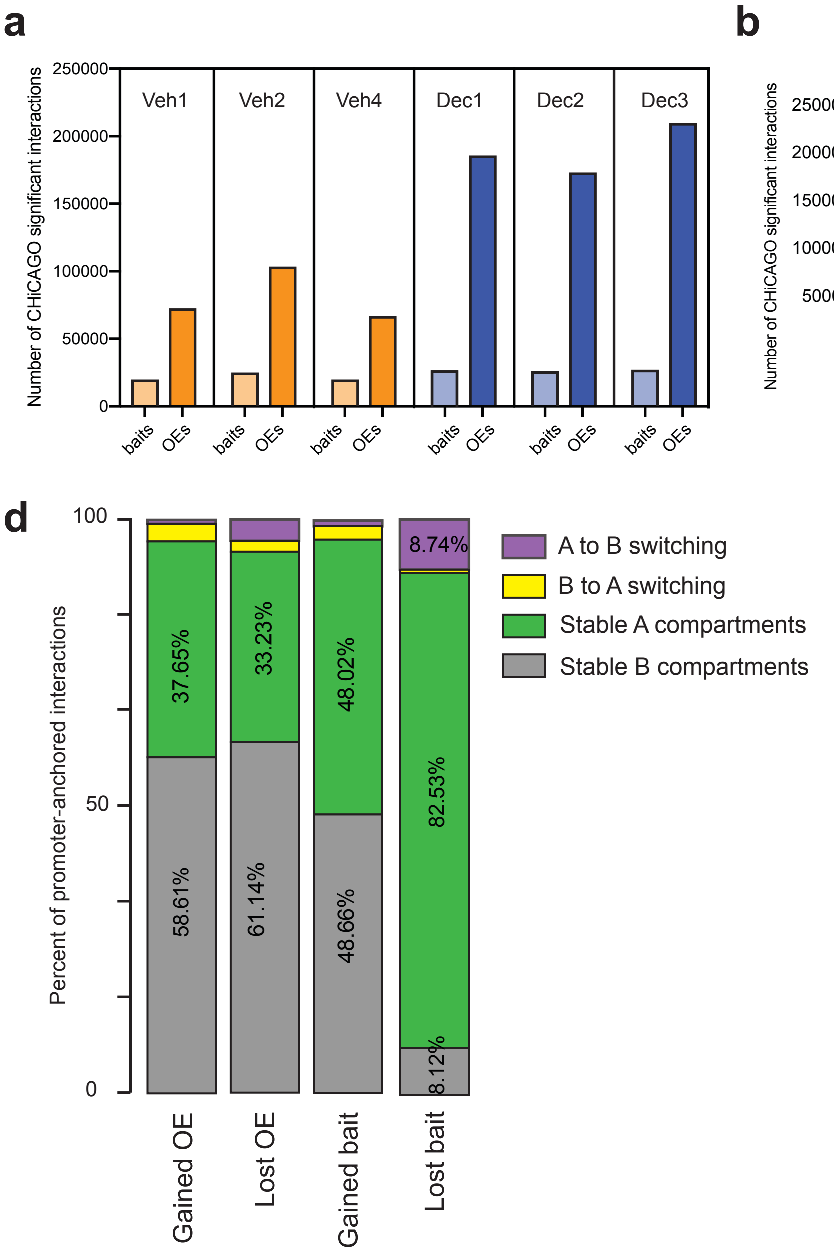

**Extended Data Fig. 3**

**(a)** Number of promoter baits and enhancer OEs involved in significant CHiCAGO interactions for each of the PCHi-C maps from Vehicle and Decitabine-treated Gar15-13 tumours (n = 3 each).

**(b)** Average number of promoter baits and enhancer OEs involved in significant CHiCAGO interactions across the three Vehicle and three Decitabine-treated PCHi-C replicates. Error bars indicate SD. *P* value Wilcoxon rank-sum test.

**(c)** Boxplot of ChICAGO scores of promoter-anchored interactions identified from PCHi-C data in Decitabine and Vehicle tumours. Data from n = 3 tumours shown.

**(d)** Bar plot showing the percentage of promoter-anchored interactions located in chromosomal compartments that switched assignments (A to B or B to A) and stable A or B compartments.

### Extended Data Fig. 4

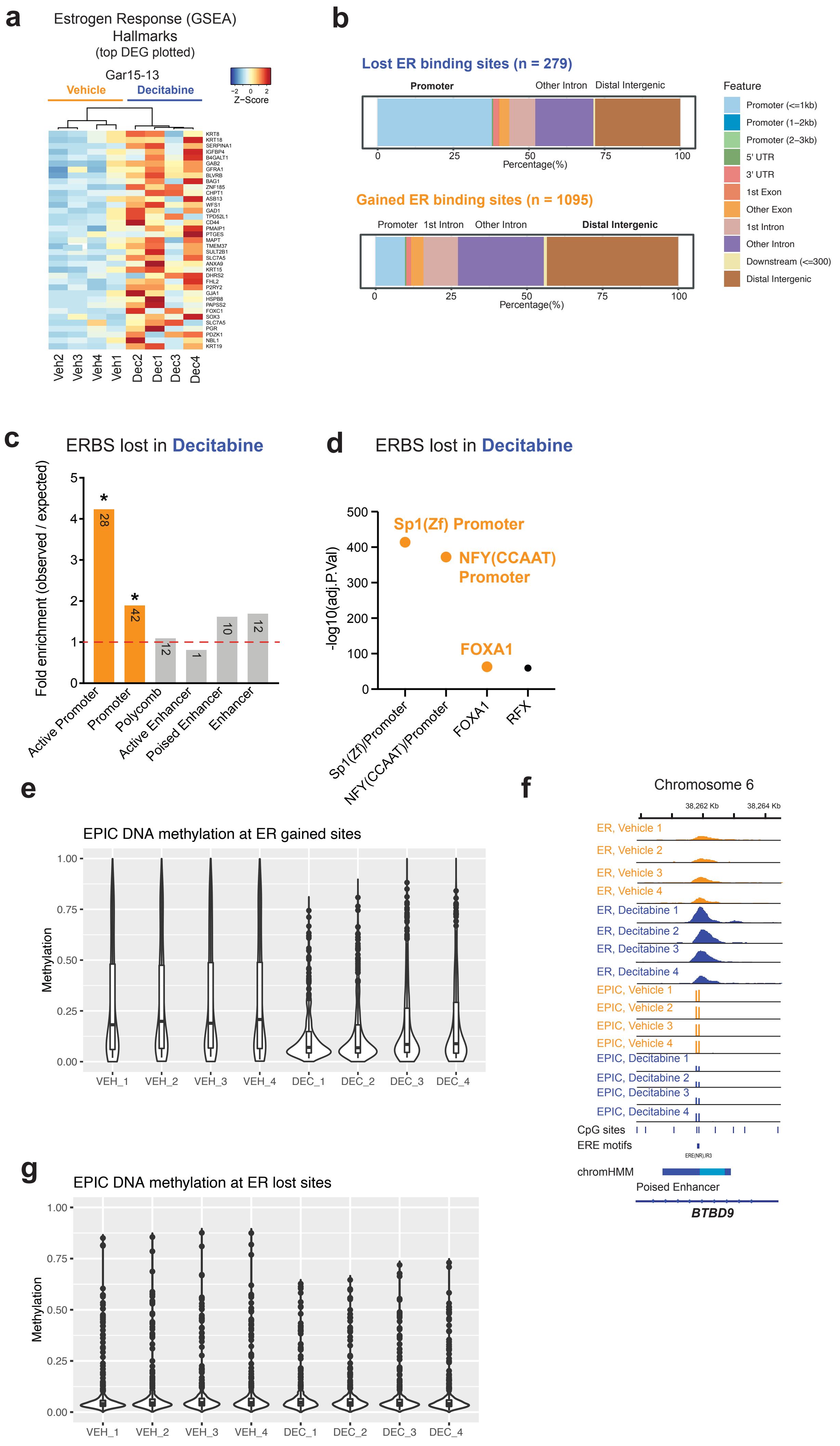

**Extended Data Fig. 4**

**(a)** RNA-seq heatmap of Decitabine-induced changes in expression of genes belonging to the Estrogen Response (GSEA) Hallmarks. Top differentially expressed genes plotted (FDR < 0.05).

**(b)** RefSeq annotation of Vehicle vs. Decitabine lost and gained ER binding sites in Gar15-13.

**(c)** ChromHMM (TAMR) annotation (**P* value < 0.001) of ER binding sites lost with Decitabine treatment compared to matched random regions across the genome. Size of the overlap is presented in the respective column.

**(d)** Transcriptions factor motifs enriched at ER binding sites lost with Decitabine treatment compared to matched random regions generated from ERE binding motifs across the genome.

**(e)** Violin plot showing DNA methylation levels at gained ER binding sites in Decitabine-treated (n = 4) and Vehicle (n = 4) PDX Gar15-13 tumours.

**(f)** Browser snapshot of ER ChIP-seq and EPIC DNA methylation (Vehicle and Decitabine-treated tumours, n = 4 each), showing concomitant gain of ER binding and loss of DNA methylation at enhancer of ER target gene *BTBD9*.

**(g)** Violin plot showing DNA methylation levels at lost ER binding sites in Decitabine-treated (n = 4) and Vehicle (n = 4) Gar15-13 tumours.

### Extended Data Fig. 5

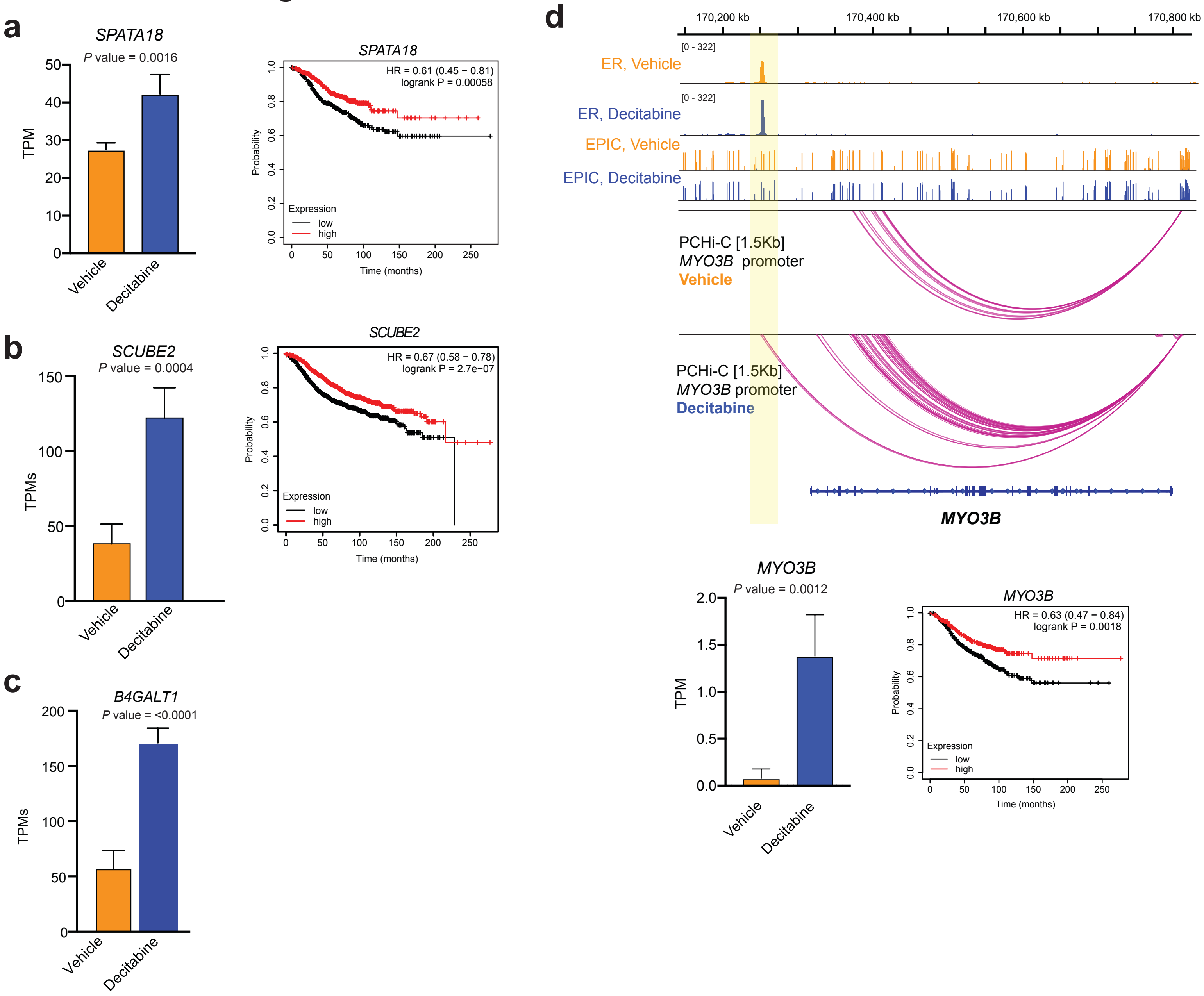

**Extended Data Fig. 5**

**(a)** The relative mRNA expression levels of *SPATA18* gene from RNA-seq (two-tailed t-test $P < 0.05$ derived from four replicates). Error bars indicate standard deviation from four samples. Kaplan–Meier survival plot showing the ability of *SPATA18* gene to stratify ER+ breast cancer patients in the METABRIC cohort into good and poor outcome groups. Data were analysed using the log-rank test. *P* values indicated within the graph.

**(b)** The relative mRNA expression levels of *SCUBE2* gene from RNA-seq (two-tailed t-test $P < 0.05$ derived from four replicates). Error bars indicate standard deviation from four samples. Kaplan–Meier survival plot showing the ability of *SCUBE2* gene to stratify ER+ breast cancer patients in the METABRIC cohort into good and poor outcome groups. Data were analysed using the log-rank test. *P* values indicated within the graph.

**(c)** The relative mRNA expression levels of *B4GALT1* gene from RNA-seq (two-tailed t-test $P < 0.05$ derived from four replicates). Error bars indicate standard deviation from four samples.

**(d)** Browser snapshots showing the promoter-anchored interactions at the *MYO3B* ER target gene, together with ER ChIP-seq, EPIC DNA methylation and ChromHMM track. Merged replicate data shown (n = 4 each). In Decitabine-treated tumours, the *MYO3B* promoter displays increased number of interactions with an enhancer, which gains ER binding with Decitabine treatment. The relative expression of the *MYO3B* gene was significantly upregulated in Decitabine-treated tumours (two-tailed t-test $P < 0.05$ derived from four replicates) and associated with good outcome in ER+ breast cancer patients in the METABRIC cohort. Data were analysed using the log-rank test. *P* values indicated within the graph.

#### Extended Data Fig. 6

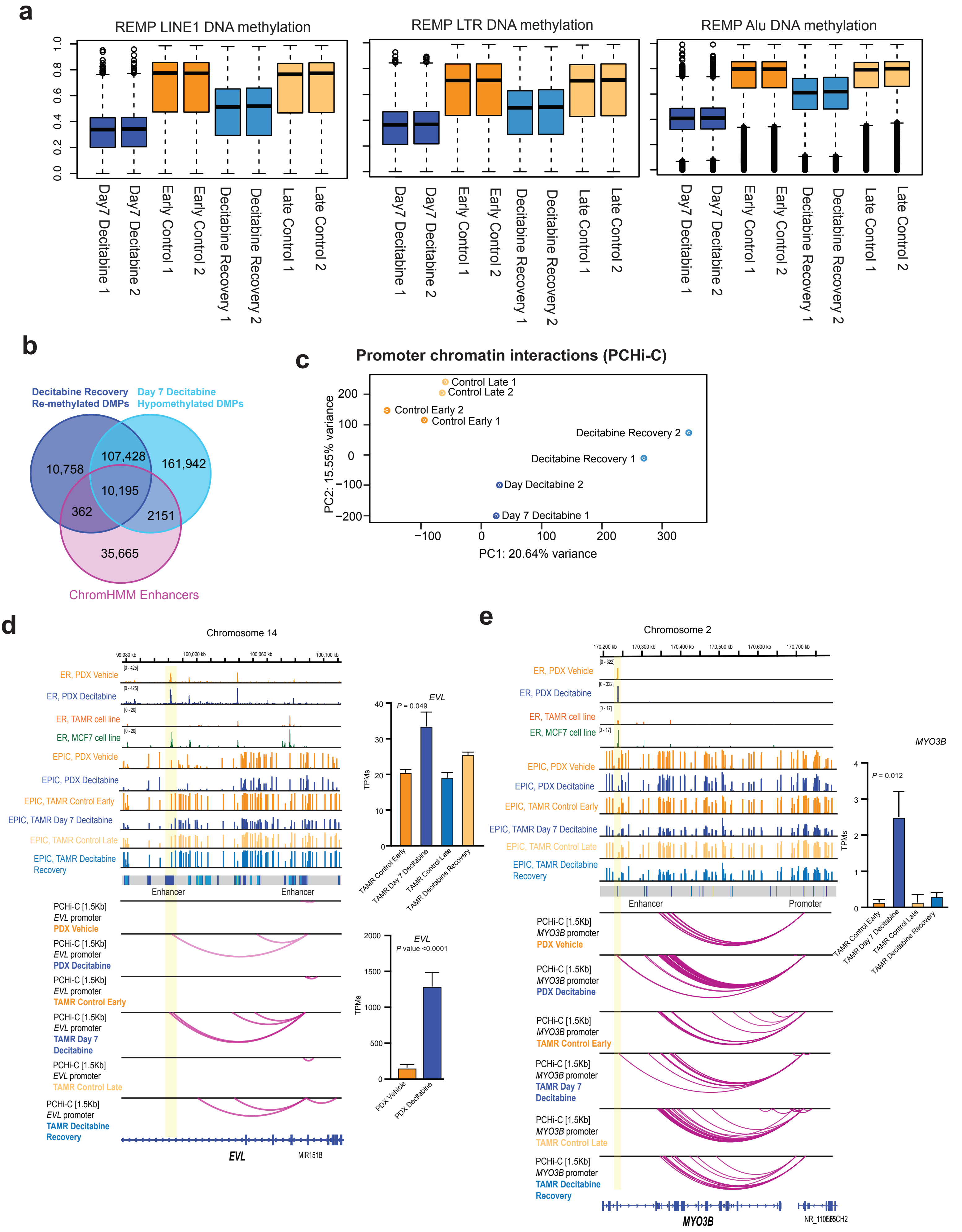

**f**

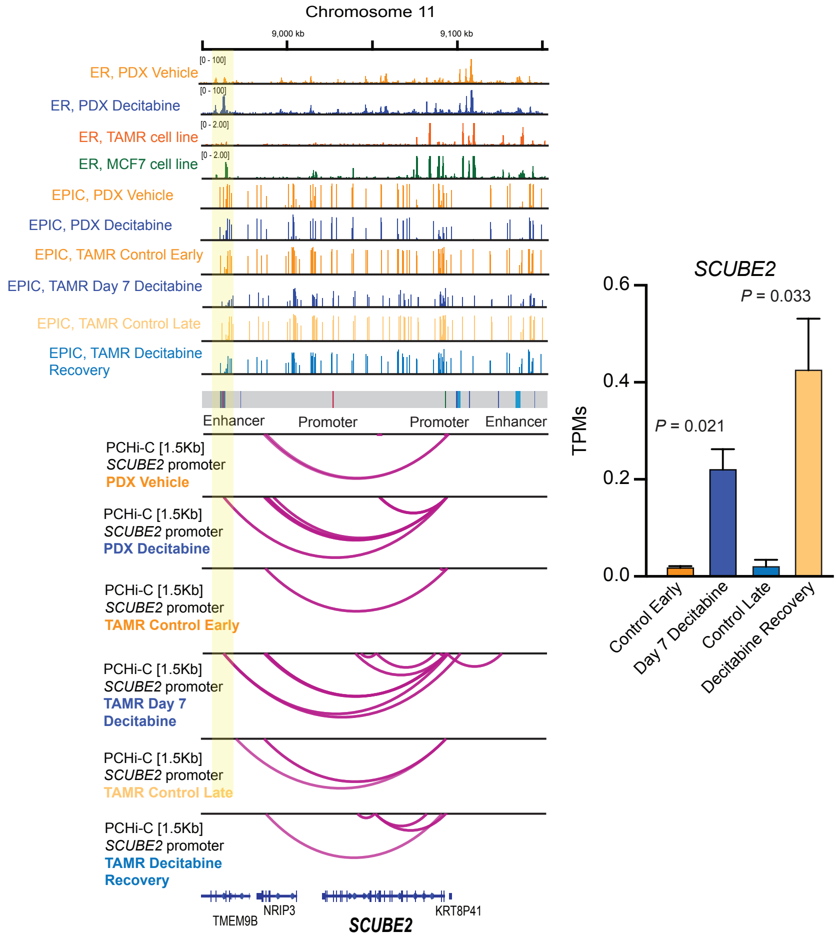

Chromosome 11
9,000 kb
9,100 kb
ER, PDX Vehicle
ER, PDX Decitabine
ER, TAMR cell line
ER, MCF7 cell line
EPIC, PDX Vehicle
EPIC, PDX Decitabine
EPIC, TAMR Control Early
EPIC, TAMR Day 7 Decitabine
EPIC, TAMR Control Late
EPIC, TAMR Decitabine Recovery
Enhancer
Promoter
Promoter
Enhancer
PCHi-C [1.5Kb] SCUBE2 promoter PDX Vehicle
PCHi-C [1.5Kb] SCUBE2 promoter PDX Decitabine
PCHi-C [1.5Kb] SCUBE2 promoter TAMR Control Early
PCHi-C [1.5Kb] SCUBE2 promoter TAMR Day 7 Decitabine
PCHi-C [1.5Kb] SCUBE2 promoter TAMR Control Late
PCHi-C [1.5Kb] SCUBE2 promoter TAMR Decitabine Recovery
TMEM9B
NRIP3
SCUBE2
KRT8P41
SCUBE2
P = 0.033
P = 0.021
TPMs
0.6
0.4
0.2
0.0
Control Early
Day 7 Decitabine
Control Late
Decitabine Recovery

**Extended Data Fig. 6**

**(a)** Boxplots showing the distribution of DNA methylation profiles for Control (Control Early and Control Late; n = 2 replicates), Decitabine-treated (Day 7 Decitabine; n = 2) and 28 days post-Decitabine treatment (Decitabine Recovery; n = 2) TAMR cells for EPIC probes mapping to LTR, LINE1 and Alu repetitive elements (REMP annotation). Black line indicates median ± SD.

**(b)** Three-way Venn diagram showing overlap of DMPs that are re-methylated in Decitabine Recovery compared to Day 7 Decitabine, Day 7 Decitabine hypomethylated DMPs and ChromHMM enhancer regions.

**(c)** Principal component analysis showing the relationship among the PCHi-C promoter-anchored interactions of TAMR cells treated with Decitabine (Day 7 Decitabine) and following 28 days recovery (Decitabine Recovery) as well as matched control cells (Control Early and Late) (n = 2). Data plotted for normalised ChICAGO interaction scores.

**(d)** Browser snapshots showing promoter-anchored interactions at the *EVL* ER target gene. Data from Gar15-13 PDX model (ER ChIP-seq, EPIC and PCHi-C) has been overlayed with TAMR cell line data (EPIC and PCHi-C) together with ER ChIP-seq for MCF7/TAMR cell lines (Ross-Innes *et al.,* 2012) and ChromHMM track. Merged replicate data shown (n = 4 each for Gar15-13 and n = 2 for TAMRs). In Decitabine-treated PDX tumours and TAMRs (Day 7 Decitabine), the *EVL* promoter displays additional interactions with intra-genic enhancer region, which gains ER binding with Decitabine treatment in PDXs, concomitant with loss of DNA methylation in both PDXs and TAMRs. These ectopic chromatin interactions are lost after 28 days of recovery with recovery of DNA methylation at that enhancer locus. Expression of the *EVL* gene was significantly upregulated in Decitabine-treated *vs.* Vehicle PDXs. In TAMRs *EVL* expression increased in Day 7 Decitabine-treated TAMRs compared to Control Early samples and was restored in Decitabine Recovery TAMRs.

**(e)** Browser snapshots showing promoter-anchored interactions at the *MYO3B* ER target gene. Data from Gar15-13 PDX model (ER ChIP-seq, EPIC and PCHi-C) has been overlayed with TAMR cell line data (EPIC and PCHi-C) together with ER ChIP-seq for MCF7/TAMR cell lines (Ross-Innes *et al.,* 2012) and ChromHMM track. In Decitabine-treated PDX tumours and TAMRs (Day 7 Decitabine), the *MYO3B* promoter displays increased number of interactions with an enhancer, which gains ER binding with Decitabine treatment in PDXs. The relative expression of the *MYO3B* gene was significantly upregulated in Decitabine-treated PDXs and TAMR cells (two-tailed t-test $P < 0.05$ derived from four replicates).

**(f)** Browser snapshots showing promoter-anchored interactions at the *SCUBE2* ER target gene. Data from Gar15-13 PDX model (ER ChIP-seq, EPIC and PCHi-C) has been overlayed with TAMR cell line data (EPIC and PCHi-C) together with ER ChIP-seq for MCF7/TAMR cell lines (Ross-Innes *et al.,* 2012) and ChromHMM track. Merged replicate data shown (n = 4 each for Gar15-13 and n = 2 for TAMRs). In Day 7 Decitabine-treated TAMRs, the *SCUBE2* promoter displays additional interactions with a distal enhancer, which gains ER binding with Decitabine treatment. Expression of the *SCUBE2* gene was significantly upregulated in Day 7 Decitabine-treated TAMRs and its expression continued to increase in Decitabine Recovery TAMRs.
